# The influence of extrachromosomal elements in the anthrax “cross-over” strain *Bacillus cereus* G9241

**DOI:** 10.1101/2022.07.13.499895

**Authors:** Grace Taylor-Joyce, Shathviga Manoharan, Thomas Brooker, Carmen Sara Hernandez-Rodrıguez, Les Baillie, Petra C. F. Oyston, Alexia Hapeshi, Nicholas R. Waterfield

## Abstract

*Bacillus cereus* G9241 was isolated from a welder who survived a pulmonary anthrax-like disease. Strain G9241 carries two virulence plasmids, pBCX01 and pBC210, as well as an extrachromosomal prophage, pBFH_1. pBCX01 has 99.6% sequence identity to pXO1 carried by *Bacillus anthracis* and encodes the tripartite anthrax toxin genes and *atxA*, a mammalian virulence transcriptional regulator. This work looks at how the presence of pBCX01 and temperature may affect the lifestyle of *B. cereus* G9241 using a transcriptomic analysis and by studying spore formation, an important part of the *B. anthracis* lifecycle. Here we report that pBCX01 has a stronger effect on gene transcription at the mammalian infection relevant temperature of 37°C in comparison to 25°C. At 37°C, the presence of pBCX01 appears to have a negative effect on genes involved in cell metabolism, including biosynthesis of amino acids, whilst positively affecting the transcription of many transmembrane proteins. The study of spore formation showed *B. cereus* G9241 sporulated rapidly in comparison to the *B. cereus* sensu stricto type strain ATCC 14579, particularly at 37°C. The carriage of pBCX01 did not affect this phenotype suggesting that other genetic elements were driving rapid sporulation. An unexpected finding of this study was that pBFH_1 is highly expressed at 37°C in comparison to 25°C and pBFH_1 expression leads to the production of Siphoviridae-like phage particles in the supernatant of *B. cereus* G9241. This study provides an insight on how the extrachromosomal genetic elements in *B. cereus* G9241 has an influence in bacterial phenotypes.

## INTRODUCTION

The *Bacillus cereus* sensu lato group contains bacterial species from diverse ecological niches and life styles (Jensen *et al*., 2003). All members of the group, which includes *Bacillus anthracis, Bacillus cereus* and *Bacillus thuringiensis* among other ecologically diverse species, have a similar chromosomal genetic composition. It is clear that the distinct phenotypes and life cycles of various species are primarily defined by the different extrachromosomal elements they carry (Rasko *et al*., 2005). The insect pathogenic *B. thuringiensis* strains encode insecticidal toxins, such as the delta endotoxins, on plasmids such as the large pBtoxis plasmid (Stein *et al*., 2006). Members of the *B. cereus* sensu stricto species can carry a range of different plasmids that define the severity of opportunistic human infections they can cause. A good example being the pCER270 plasmid which encodes the cereulide toxin biosynthesis gene cluster (Rasko *et al*., 2007), responsible for severe food poisoning. In the case of *B. anthracis*, the causative agent of anthrax, carriage of the anthrax toxin plasmid pXO1 and the capsule plasmid pXO2 facilitate a lethal mammalian infection life cycle. Interestingly, all *B. anthracis* strains have a frame shift mutation in the gene for the quorum sensing global regulator, PlcR, which controls the secretion of toxins and enzymes in other members of the *B. cereus* sensu lato group (Agaisse *et al*., 1999).

*B. cereus* strain G9241 (from here on referred to as *Bc*G9241) was previously identified as the causative agent of an anthrax-like pneumonia infection of a metal worker in Louisiana in 1994. Phenotypic and whole genome sequencing analyses confirmed that the isolate was a *B. cereus* sensu stricto species which carries a close homologue of the *B. anthracis* plasmid pXO1, named pBCX01, encoding the tripartite anthrax toxins and their mammalian host responsive regulator AtxA1 (Hoffmaster *et al*., 2004). *Bc*G9241 also carries a plasmid named pBC210 which, while unrelated to the *B. anthracis* plasmid pXO2, also encodes capsule genes, albeit distinct from those on pXO2 and a second *atxA1* like regulator gene designated *atxA2*. The third extrachromosomal element found in *Bc*G9241 is pBFH_1, a linear phagemid (Oh *et al*., 2011). Although the sequence is available for pBFH_1, it is not known if/how it contributes to the lifestyle of *Bc*G9241. While *Bc*G9241 carries these three plasmids and produces anthrax toxins, *Bc*G9241 is resistant to γ-phage and penicillin and carries an intact copy of the *plcR* gene, hence why it is referred to as a *B. cereus* strain (Hoffmaster *et al*., 2004). A small number of other *B. cereus* strains that are capable of causing an anthrax-like disease have also been isolated and are collectively referred to as ‘cross-over’ strains, an in-depth description of these strains given in a recent review (Baldwin, 2020). Since this review was published, two further ‘cross-over’ strains have been isolated from patients with an anthrax like disease (Dawson *et al*., 2021).

*B. cereus* and *B. anthracis* differ from each other in the disease states they cause and also have very different lifestyles. *B. cereus* is an opportunistic pathogen causing primarily gastrointestinal infections in humans. *B. cereus* is also often found in the guts of many invertebrates, which both facilitates its dissemination in the environment and provides a niche for a potential necromonic lifestyle (Margulis *et al*., 1998; Swiecicka and Mahillon, 2006). In contrast, *B. anthracis* is primarily adapted to cause acute mammalian infections abandoning the more generalist lifestyle of its recent *B. cereus* ancestor. *B. anthracis* infects its mammalian host as a spore, often picked up from the soil by grazing livestock or wildlife through inhalation or ingestion (Bellan *et al*., 2013). The animal then dies from the acute anthrax disease. Vegetative *B. anthracis* cells present in the carcass sporulate as nutrients become limited and re-enter the soil as a spore, completing the life cycle (Lindeque and Turnbull, 1994). This life cycle relies on *B. anthracis* being able to successfully kill its host and sporulate once its host is deceased. Current studies of *Bc*G9241 have focused on the contribution of the anthrax toxins and the tetrasaccharide and hyaluronic acid capsules to the ability of *Bc*G9241 to cause a fatal anthrax-like disease (Scarff *et al*., 2018). The contribution of chromosomal regulation by pBCX01 or AtxA1 to virulence is less well understood and the sporulation of *Bc*G9241 has so far not been investigated.

This study looks at how closely life cycle related phenotypic traits of *Bc*G9241 resemble those of *B. anthracis* and how pBCX01 may influence this. To do this we have carried out RNAseq experiments to elucidate how carriage of pBCX01 influences the transcriptome of *Bc*G9241 at the mammalian relevant temperature of 37 °C compared to the more environmentally appropriate temperature of 25 °C. We have also assessed the sporulation phenotype of *Bc*G9241. Sporulation is an essential part of the *B. anthracis* life cycle and is intrinsically linked to other cellular processes, including virulence, through a complex regulatory network (Tan and Ramamurthi, 2014). The results presented here, as well as other work done by our group (Manoharan *et al*., 2022), show that *Bc*G9241 phenotypes have a strong temperature dependence allowing it to phenocopy *B. anthracis-*like behavior at 37 °C, while conversely behaving like *B. cereus* sensu stricto at 25 °C. pBCX01 has a strong influence on the transcriptome at 37 °C, including the suppression of the transcription of genes involved in metabolic and biosynthetic pathways whilst simultaneously upregulating the expression of genes involved in the import of nutrients, an effect which is not seen at 25 °C. We propose that this may facilitate mammalian virulence, allowing *Bc*G9241 to have a growth advantage whilst simultaneously restricting nutrients available to the host. In addition, an unexpected finding of our study showed a significant increase in the pBFH_1 lysogenic phagemid expression at 37 °C compared to 25 °C. Interestingly, close homologues of this phagemid are also carried by other ‘cross-over’ strains, despite no direct phylogenetic relationships between the chromosomes, suggesting a common role in their life cycles. We further demonstrated that pBFH_1 produces Siphoviridae-like phage particles in the supernatant of *Bc*G9241. Our observations regarding sporulation show that *Bc*G9241 has a temperature dependent sporulation phenotype and sporulates rapidly compared to *B. cereus* ATCC 14579 (from here on referred to as *Bc*ATCC 14579), particularly at 37 °C, which is independent of the presence of pBCX01.

In summary, our results show that pBCX01 has allowed *Bc*G9241 to adopt a life cycle that resembles that of *B. anthracis* and that the response of *Bc*G9241 to temperature is crucial to this. We have also demonstrated that other genetic elements contribute to this temperature dependent phenotype independently of pBCX01. To allow us to examine the influence of pBCX01 on bacterial phenotypes, we constructed a plasmid cured strain designated *Bc*G9241 ΔpBCX01.

## RESULTS

### The influence of pBCX01 on global *Bc*G9241 transcription during exponential growth

To determine if pBCX01 influences BcG9241 at a transcriptional level, mRNA was extracted from cultures of *Bc*G9241 WT (WT) and *Bc*G9241 ΔpBCX01 (ΔpBCX01) growing exponentially (OD_600_=0.5) at both 25 °C and 37 °C. An RNAseq analysis was performed to compare the transcriptomes of WT and ΔpBCX01. A comparison of the transcriptomes of each strain at 25 °C and 37 °C was also performed. The full datasets generated can be seen in the **Supplementary Datasets 1-4**.

### The transcriptional response of *Bc*G9241 to temperature is influenced by pBCX01

The number of genes found to be differentially expressed between 37 °C and 25 °C during exponential growth of the WT strain was 250 (Figure 1A, shown in green), while for the ΔpBCX01 strain the number was considerably higher, exhibiting 1,184 differentially expressed genes (DEGs) (Figure 1A, shown in orange). Furthermore, there were only 177 genes that are differentially expressed between 25 °C and 37 °C shared between WT and ΔpBCX01 (Figure 1A). This demonstrates that the pBCX01 plasmid has a potent influence on temperature dependent expression of the chromosomal genes. To compare the functions of temperature dependent DEGs of WT and ΔpBCX01, we performed gene enrichment analyses using the STRING online tool (Szklarczyk *et al*., 2019) and focused on the KEGG pathways that were found to be enriched. Unfortunately, this database does not recognize the *Bc*G9241 locus tags, therefore all genes of the *Bc*G9241 genome were assigned a *B. anthracis* Sterne locus tag, in cases where there was amino acid homology of 90% or more. As such, *Bc*G9241 genes that had no homologue in the *B. anthracis* genome have been excluded in this analysis.

**Figure 1.**
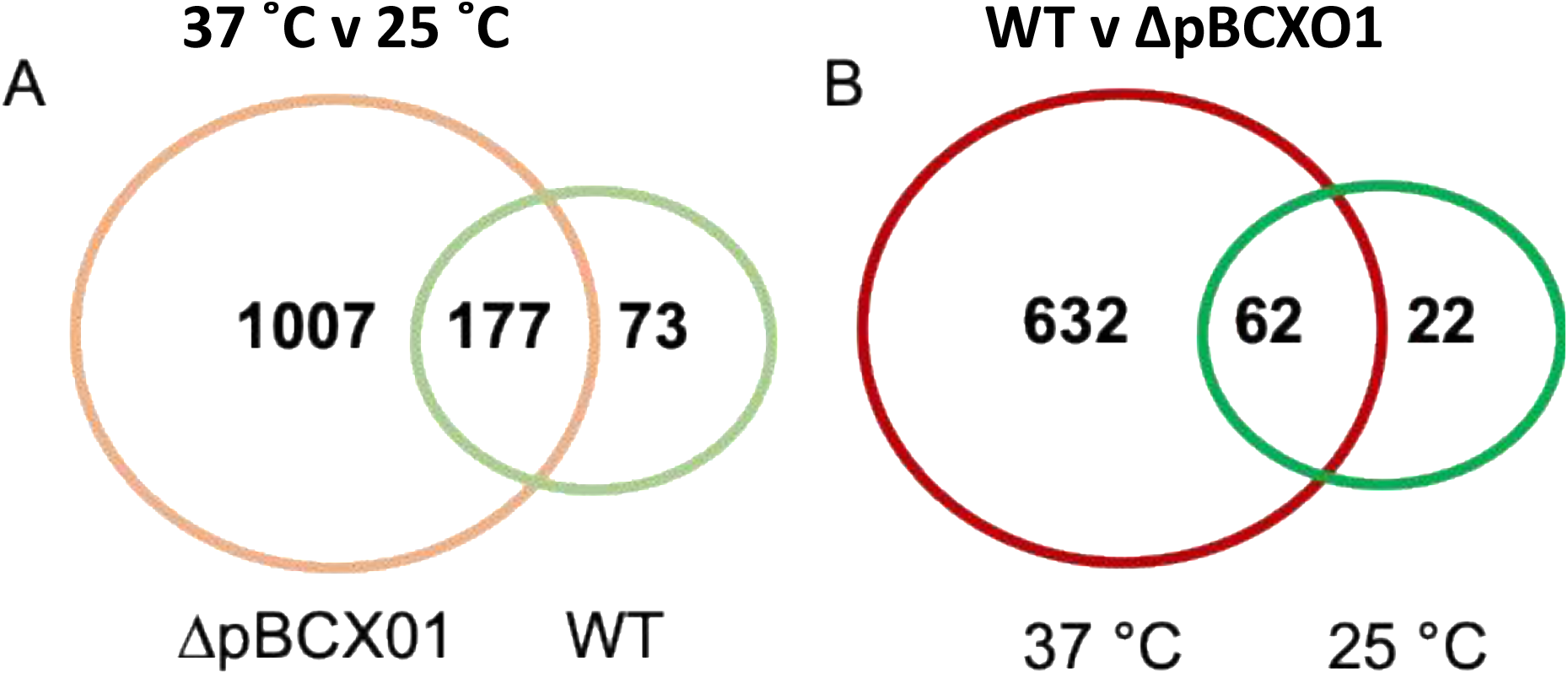
Venn diagrams of genes that are differentially expressed between **A)** 37° C and 25° C in *Bc*G9241 WT (green) and BcG9241 ΔpBCX01 (orange) and **B**) *Bc*G9241 WT and ΔpBCX01 when cultured at either 25° C (green) or 37° C (red).

In ΔpBCX01, genes significantly differentially expressed between 37 °C and 25 °C showed a significant enrichment for genes involved in KEGG pathways associated with an increased growth rate. These include genes involved in pathways relating to metabolism, the ribosome and biosynthesis of secondary metabolites and amino acids (Table 1). The majority of these DEGs had a higher relative transcript abundance at 37 °C compared to 25 °C which is consistent with *Bc*G9241 having an increased protein production and higher growth rate at 37 °C compared to 25 °C (Figure S1). In contrast, the genes with temperature dependent differential expression in the WT strain did not show the same profile of functions associated with an increased growth rate. In the WT there were 16 genes involved in metabolic pathways that had a higher relative transcript abundance at 37 °C compared to 25 °C but surprisingly, no genes involved in amino acid biosynthesis or ribosomal genes. This is unexpected as both strains exhibit an increased growth rate at 37 °C in comparison to that at 25 °C (Figure S1). An analysis of the genes that had a higher relative transcript abundance at 25 °C compared to 37 °C in the WT showed there was a significant enrichment for genes involved in bacterial chemotaxis. Five of the six of these differentially expressed bacterial chemotaxis genes were not found to be differentially expressed in ΔpBCX01.

**Table 1:** KEGG pathway enrichment analysis results of genes with a significantly higher transcript abundance at 37°C relative to 25°C at exponential phase in *Bc*G9241 WT and *Bc*G9241 ΔpBCX01. A false discovery rate cut off 0.01 was used to accept a significant enrichment of DEG in each group.

| KEGG pathway | Observed gene count | No. of genes in pathway | False discovery rate |
| --- | --- | --- | --- |
| *Bc* G9241 WT, 37°C > 25°C |  |  |  |
| Chloroalkane and chloroalkene degradation | 3 | 11 | 0.0095 |
| Glycolysis / Gluconeogenesis | 5 | 50 | 0.0095 |
| Pyruvate metabolism | 5 | 55 | 0.0095 |
| Microbial metabolism in diverse environments | 9 | 216 | 0.0095 |
| Metabolic pathways | 16 | 657 | 0.0095 |
| *Bc* G9241 ΔpBCX01, 37°C > 25°C | | | |
| Metabolic pathways | 161 | 657 | 1.26E-29 |
| Biosynthesis of secondary metabolites | 92 | 320 | 1.89E-19 |
| Biosynthesis of antibiotics | 76 | 235 | 2.18E-18 |
| Microbial metabolism in diverse environments | 54 | 216 | 3.22E-09 |
| Ribosome | 25 | 56 | 3.30E-08 |
| Carbon metabolism | 36 | 117 | 3.37E-08 |
| Biosynthesis of amino acids | 39 | 140 | 6.44E-08 |
| Pyrimidine metabolism | 21 | 56 | 4.45E-06 |
| Glycolysis / Gluconeogenesis | 19 | 50 | 1.21E-05 |
| Purine metabolism | 21 | 68 | 4.63E-05 |
| Aminoacyl-tRNA biosynthesis | 14 | 35 | 0.00016 |
| Citrate cycle (TCA cycle) | 12 | 27 | 0.00026 |
| Propanoate metabolism | 13 | 33 | 0.00031 |
| Oxidative phosphorylation | 15 | 46 | 0.00045 |
| Pyruvate metabolism | 16 | 55 | 0.00074 |

### At 37 °C pBCX01 has a much greater influence on the transcriptome than at 25 °C

To look at the direct effect of pBCX01 on the transcriptome at both 37 °C and 25 °C a list of genes that were differentially expressed between WT and ΔpBCX01 was generated for cultures grown at the two temperatures. The number of genes found to be differentially expressed between WT and ΔpBCX01 when cultures were grown at 37 °C was 694 (Figure 1B, shown in red), whereas when cultures were grown at 25 °C the number of DEG was lower at 84 (Figure 1B, shown in green). Of the 84 genes differentially expressed between WT and ΔpBCX01 at 25 °C, 62 of them were also differentially expressed at 37 °C between WT and ΔpBCX01. In addition to this, many of the genes with the highest differential expression between WT and ΔpBCX01 at 25 °C are also seen to be differentially expressed at 37 °C (Table S1-S2). This indicates that at 37 °C pBCX01 has a greater influence on transcription than it does at 25 °C.

### pBCX01 suppresses metabolic and biosynthetic processes at 37 °C, while conversely having a positive effect on the transcription of many transmembrane proteins

To take a closer look at the DEGs between WT and ΔpBCX01 at 37 °C the list of DEGs was split into those that had a higher relative transcript abundance in either the WT or ΔpBCX01. As before, STRING was used to perform an enrichment analysis for each list of DEGs. Genes with higher relative transcript abundance in WT compared to ΔpBCX01 at 37 °C had a significant enrichment of genes encoding transmembrane proteins (Table 2). Many of these transmembrane proteins have predicted transport functions, including transporters of cobalt, formate/nitrite, phosphonates and amino acids. Four of the top 15 most differentially expressed genes showing higher transcription in WT have a predicted transporter function (Table S1). One of these encoded an EcsB family protein involved in regulation and secretion of extracellular proteins. A gene encoding InhA1 also had a higher transcription level in WT compared to ΔpBCX01 at 37 °C. InhA1 is a metalloprotease involved in breaking down host proteins for nutrients (Terwilliger *et al*., 2015) as well as mediating escape of *B. cereus* from macrophage (Haydar *et al*., 2018). Interestingly, pBCX01 also has a positive effect on genes predicted to encode transcriptional regulators as well as the gene encoding the transition state regulator AbrB.

**Table 2:** KEGG pathway enrichment analysis results of genes significantly affected by the presence of pBCX01 at 37°C exponential phase. A false discovery rate cut off or 0.01 was used to accept a significant enrichment of DEG in each group.

| KEGG pathway | Observed gene count | No. of genes in pathway | False discovery rate |
| --- | --- | --- | --- |
| Positively affected by the presence of pBCX01 |  |  |  |
| Transmembrane | 98 | 984 | 3.02E-08 |
| Transmembrane helix | 96 | 973 | 4.14E-08 |
| Negatively affected by the presence of pBCX01 |  |  |  |
| Metabolic pathways | 72 | 657 | 4.14E-12 |
| Biosynthesis of antibiotics | 41 | 235 | 4.51E-12 |
| Biosynthesis of secondary metabolites | 47 | 320 | 1.09E-11 |
| Carbon metabolism | 22 | 117 | 5.36E-07 |
| Biosynthesis of amino acids | 23 | 140 | 1.85E-06 |
| Propanoate metabolism | 10 | 33 | 7.42E-05 |
| Microbial metabolism in diverse environments | 25 | 216 | 0.00013 |
| Fatty acid metabolism | 9 | 32 | 0.00027 |
| Valine, leucine and isoleucine degradation | 7 | 21 | 0.00076 |
| Butanoate metabolism | 8 | 29 | 0.00076 |
| Fructose and mannose metabolism | 5 | 16 | 0.0086 |
| Terpenoid backbone biosynthesis | 5 | 16 | 0.0086 |

The analysis of genes with higher relative transcript abundance in ΔpBCX01 compared to WT showed that there was a significant enrichment for genes involved in metabolism, biosynthesis of secondary metabolites and amino acids. There is a strong similarity between the DEGs seen between 37 °C and 25 °C in ΔpBCX01 and the DEGs with higher transcript abundance in ΔpBCX01 when grown at 37 °C. Both have significant enrichments in genes involved in metabolism, biosynthesis of secondary metabolites and amino acids. This pattern of DEGs suggests that at 37 °C pBCX01 is suppressing the expression of groups of genes involved in metabolism, biosynthesis of secondary metabolites and biosynthesis of amino acids, processes that would normally be associated with an increase in growth rate.

Interestingly, the five transcripts with the highest relative abundance in ΔpBCX01 compared to WT when grown at 37 °C were all around 110 nucleotides in length and encoded adjacent to each other in the genome (Table S1). Homologues of these five genes can be found in the same arrangement in other *B. cereus* and *B. anthracis* genomes. The five genes are all homologues of each other exhibiting a DUF3948, but with no known function.

To compare the overall transcriptional profiles for all four conditions (WT grown at 37 °C and 25 °C and ΔpBCX01 grown at 37 °C and 25 °C) a principal component analysis (PCA) plot was generated using data from each biological replicate (Figure 2). Principal component one describes 55% of the variance and principal component two describes 20% of the variance between all samples. All biological replicates generally cluster together. There is some overlap between the WT and ΔpBCX01 samples cultured at 25 °C, supporting the hypothesis that pBCX01 has a stronger influence on the transcriptome at 37 °C than at 25 °C. Interestingly, the PCA plot shows that the WT 37 °C samples cluster closer to all the 25 °C samples than they do to the ΔpBCX01 37 °C samples. This may be due to activity of pBCX01 at 37 °C and its suppressive effect on genes that would otherwise be upregulated in response to a more optimal growth temperature.

**Figure 2:**
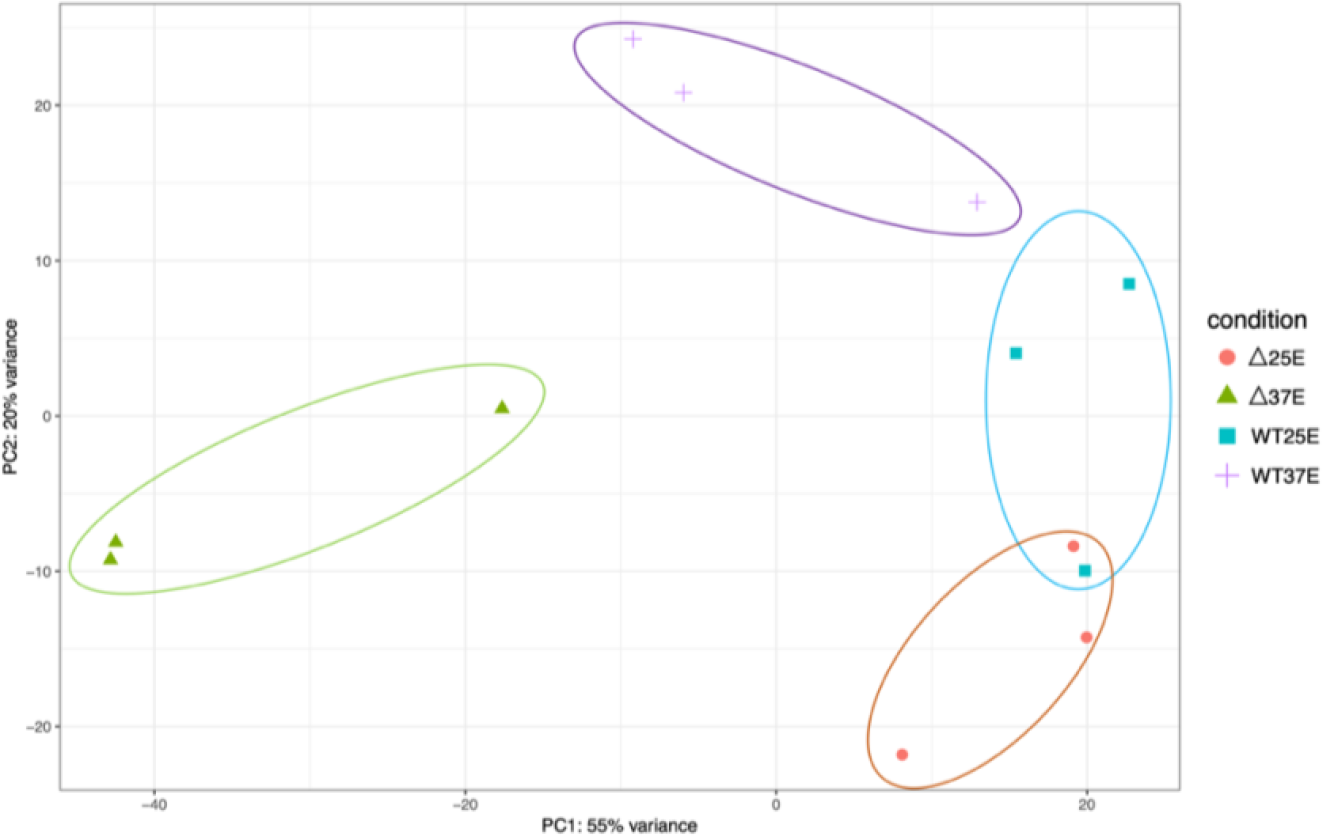
A principal component analysis generated from a DEseq2 analysis of the RNAseq data from *Bc*G9241 WT grown at either 37°C or 25°C and *Bc*G9241 ΔpBCX01 grown at either 37°C or 25°C. The x axis, showing principal component 1, describes 55% of the variance in the data and the y axis, showing principal component 2, describes 20% of the variance in the data.

### The impact of pBCX01 upon the nutritional requirements of *Bc*G9241

The results of the transcriptome analysis showing that pBCX01 had a negative effect on genes that are associated with an increase growth rate could suggest that at 37 °C, WT has a slower growth rate than ΔpBCX01. Nevertheless, this does not appear to be the case when grown in nutrient rich LB, although at 37 °C the growth rate of ΔpBCX01 is slightly higher than WT, the difference is negligible (Figure S1). We therefore aimed to investigate whether pBCX01 had any effect on growth rate when cultured in a media designed to resemble the nutrient availability of the mammalian host. Additionally, we wanted to specifically assess the effect of pBCX01 on the ability of *Bc*G9241 to synthesize amino acids as many amino acid biosynthesis genes were seen to be negatively affected by pBCX01. A negative effect of AtxA on amino acid genes has been previously observed in *B. anthracis* where either the *atxA* gene had been removed, or when the bacteria were grown under conditions where AtxA is known to be less active (Bourgogne *et al*., 2003; Panda *et al*., 2014; Raynor *et al*., 2018). This strongly suggests that it may be the pBCX01 encoded AtxA1 activity which causes the observed down regulation of amino acid biosynthesis genes.

A minimal media designed to mimic the nutrient availability of the blood, referred to as blood serum mimic (BSM), was used (Terwilliger *et al*., 2015) and individual amino acids were omitted to assess the ability of WT and ΔpBCX01 to synthesize specific amino acids. *Bc*ATCC 14579 was also included in these experiments to see how the growth of WT and ΔpBCX01 compare to a typical *B. cereus* sensu stricto strain. It should be noted that for *Bc*G9241 to grow in BSM we had to supplement this medium with 0.04 μM biotin and 0.056 mM thiamine, or no growth was observed. It was anticipated that if WT exhibited a lower expression of the relevant amino acid biosynthesis genes than ΔpBCX01, then it may have a slower growth rate in media in which any of the relevant amino acids were limiting. To predict which amino acids might be most affected by the influence of pBCX01 a KEGG metabolic pathway analysis was used to construct a map of amino acid biosynthesis genes that were differentially expressed between WT and ΔpBCX01 (Figure S2). Based on this analysis; arginine, histidine, isoleucine, leucine, lysine, methionine, tryptophan and valine were selected as candidates to be individually removed from the defined growth media for testing.

Our findings indicated that both WT and ΔpBCX01 were unable to grow without leucine or valine being present in the media and that *Bc*ATCC 14579 could not grow without methionine or valine. In media with either, arginine, isoleucine, lysine or tryptophan removed or with all amino acids present, WT grew slightly better than ΔpBCX01 and although *Bc*ATCC 14579 initially grew faster, WT reached a higher OD_600_ than either *Bc*ATCC 14579 or ΔpBCX01. In media with multiple amino acids removed, *Bc*ATCC 14579 was unable to grow, presumably because of the lack of methionine. While WT grew slightly better than ΔpBCX01, there was very little growth of either strains over 24 hours. Growth curves can be seen in Figure 3.

**Figure 3:**
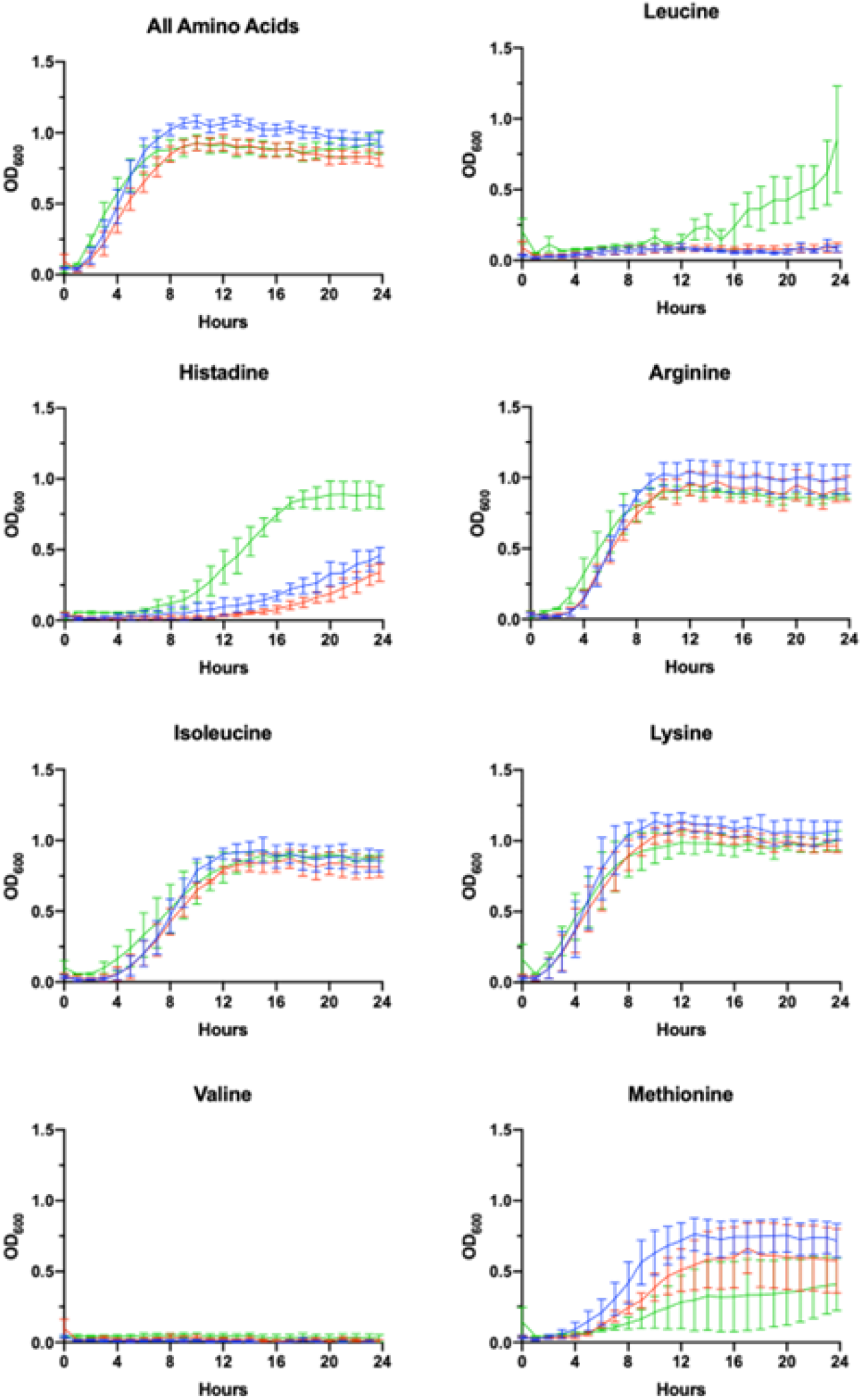
Growth in AA limiting media. Growth curves showing optical density measured at 600mn wavelength (OD_600_) plotted against time of *Bc*G9241 WT (blue line), *Bc*G9241 ΔpBCX01 (red line) and *Bc*ATCC 14579 (green line). The bars show standard error. Cultures were grown at 37°C with shaking in blood serum mimic (BSM) media with a certain amino acid omitted. The amino acid written above each graph indicates which amino acid was omitted from the growth media.

### Growth at 37°C induces pBFH_1 phagemid encoded virion production in

***Bc*G9241**

The transcriptomic comparison of both WT and ΔpBCX01 grown at 25 °C and 37 °C during exponential growth phase also revealed that the pBFH_1 phagemid genes become highly transcribed during growth at 37 °C compared to 25 °C. The most highly induced genes at 37 °C in WT included two encoding subunits of the predicted phage terminase protein, presumably involved in packing the phage genome into the maturing capsid structure. The large (AQ16_5899) and small ATPase subunit (AQ16_5898) genes were log2-fold 3.86 and 4.14 times higher at 37 °C compared to 25 °C respectively (Table S3). Transcription of 12 genes encoding capsid or tail proteins were also highly induced at 37 °C (Figure 4-red squares). Perhaps not unexpectedly a phage anti-repressor encoding gene (AQ16_5855) was seen to be induced at 37 °C. Furthermore, many transcriptional activators carried by pBFH_1 showed higher transcription at 37 °C, including two Xre superfamily like transcriptional regulators (AQ16_ 5850 and 5853). Interestingly a phagemid autolysin regulatory protein was also induced at 37 °C. Taken together this suggested that the lysogenic pBFH_1 phagemid enters a lytic cycle upon exposure to the higher temperature, as evidenced by the increased transcription of virion structural protein genes. The transcriptomic comparison of ΔpBCX01 grown at 25 °C and 37 °C also showed that the majority of pBFH_1 genes were more highly expressed at 37 °C in comparison to 25 °C. Of the 15 genes most highly expressed at 37 °C in comparison to 25 °C, 10 were pBFH_1 encoded genes (Table S5)

**Figure 4.**
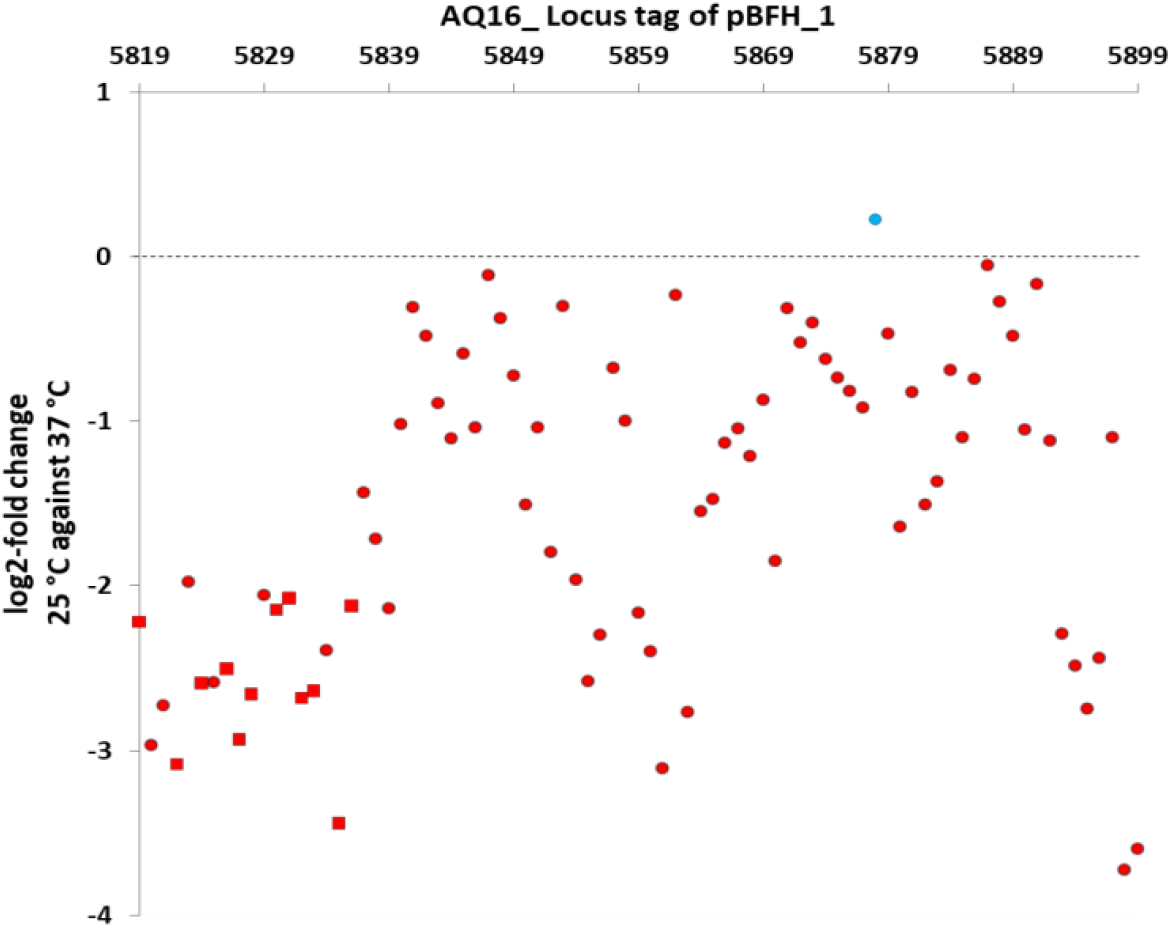
Genes on the pBFH_1 phagemid are more highly transcribed when *Bc*G9241 is grown at 37 °C, compared to at 25 °C. *Bc*G9241 was grown at 25 °C and 37 °C to exponential phase. RNA was extracted and RNA-seq performed. RNA-seq reads were processed, normalised, compared using DESEQ2 and plotted as the log2-fold change in the transcriptional level of each gene at 25 °C compared to at 37 °C. Almost all genes were more highly transcribed during growth at 37 °C (red circles/squares), with the exception of a gene encoding a Fur-regulated protein (blue circle). All known genes encoding capsid and tail proteins (red squares) were higher at 37 °C. All points plotted are significant (p value < 0.05; n=3).

A mass spectrometry proteomic analysis of supernatants of mid-exponentially phase WT culture grown at 37 °C and 25 °C was carried out by our group (Manoharan *et al*., 2022). In agreement with the transcriptomic analysis, the proteomic analysis showed that the 8 highest proteins and 40% of the total proteins higher at 37 °C compared to 25 °C were encoded on the pBFH_1 phagemid (Table 3).

**Table 3.** Supernatant proteins showing the largest increase in abundance at 37 °C compared to 25 °C during exponential growth of *Bc*G9241 WT.

| Log2-Fold Change | Protein | Gene Loci (AQ16_) |
| --- | --- | --- |
| 5.27 | (Gp49) Phage family protein | 5822 |
| 4.97 | (Gp34) Putative phage major capsid protein | 5824 |
| 4.53 | Prophage minor structural protein | 5836 |
| 4.32 | (Gp14) Putative gp14-like protein | 5832 |
| 4.32 | N-acetylmuramoyl-L-alanine amidase family protein | 5839 |
| 3.66 | Phage tail family protein | 5835 |
| 3.41 | (GpP) Putative major capsid protein | 5831 |
| 2.90 | Uncharacterized protein | 5823 |

The identification of pBFH_1 structural proteins in the supernatants of WT grown at 37 °C, led us to investigate if intact phage particles were present. Particulate material was filtered from supernatants and visualised using negative stain transmission electron microscopy. Figure 5 shows the presence of a phage with Siphoviridae like morphology. The average tail dimensions are 193 × 10 nm and the average head diameter is 60 nm which is consistent with the morphology of the Siphoviridae family (Hendrix *et al*., 2012). Like other Siphoviridae the phage particles appear to have icosahedral shaped heads and a tail made up of stacked discs. This classification is consistent with the analysis of the pBFH_1 sequence by Phaster, which assigns the structural capsid proteins as Siphoviridae (Zhou *et al*., 2011). Furthermore, PCR analysis of the phage preparations confirmed that intact phage particles carrying pBFH_1 DNA were present. Interestingly we were not able to detect phage proteins in the cellular proteome at 37 °C (Manoharan *et al*., 2022) potentially indicating that the phage had been rapidly induced and released from only a sub-population of the culture. Indeed, this is consistent with the observation that there was no reduction in growth rate or apparent lysis of the whole culture at 37 °C, and a testament to the sensitivity of the proteomic analysis.

**Figure 5.**
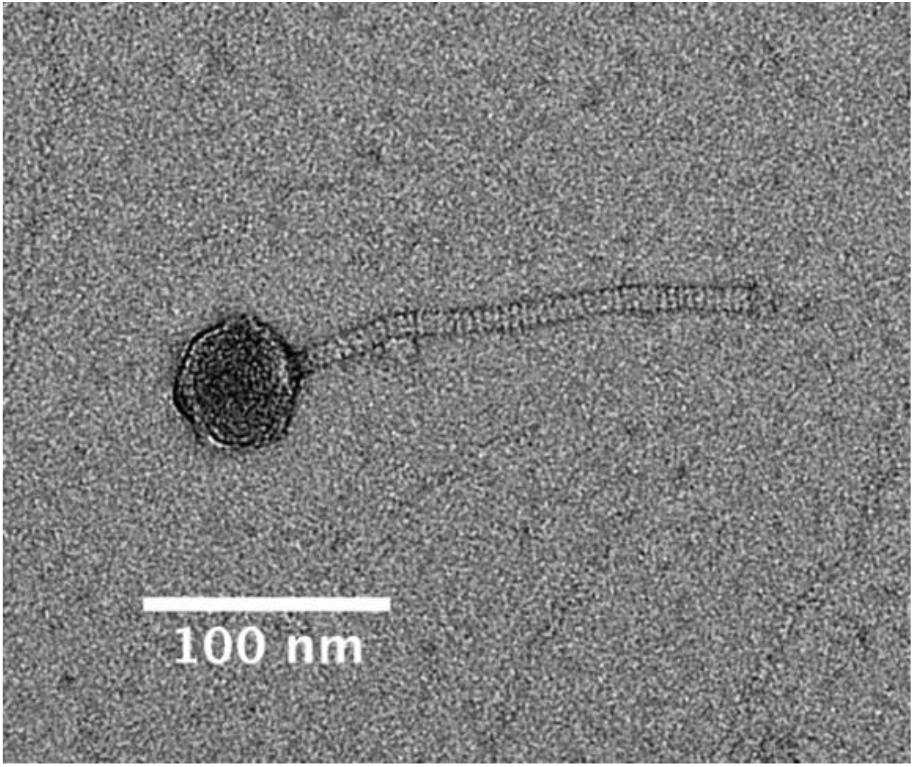
Transmission electron micrograph of the pBFH_1 phagemid virion isolated from supernatant of a *Bc*G9241 WT culture grown at 37 °C. Average tail length: 192.57 nm and average head diameter: 60.21 nm.

Nucleotide BLAST analysis was carried out to identify any other pBFH_1-like phagemids carried by other *B. cereus* or *B. anthracis* strains. All strains identified that exhibited homology to the pBFH_1 genome with a percentage coverage of 61% and above were *B. cereus* strains that were also carrying pXO1/pBCX01, pXO2 or pBC210 plasmids (Table 4). Other strains with sequences homologous to the pBFH_1 phagemid only had a sequence coverage of 43% or below in *B. cereus* strains and 17% or below in *B. anthracis* strains. There were two ‘cross-over’ strains that did not show any sequence homology to pBFH_1, 03BB102 and 03BB108. Interestingly neither 03BB102 nor 03BB108 carry pXO2 or pBC210-like plasmids, and furthermore their pBCX01 plasmids have only partial homology to pXO1.

**Table 4.** Presence of *Bc*G9241 pBFH_1 phagemid homologues in other *B. cereus* strains carrying pBCXO1 or pBCXO2. Presence or absence of pXO1/pBCX01, pXO2 and pBC210 is also indicated.

| Strain name | pXO1/pBCXO1 Present | pXO2 present | pBC210 Present | G9241 pBFH_1 homologues |  | Sample source | NCBI accession no. | Submitted by |
| --- | --- | --- | --- | --- | --- | --- | --- | --- |
| G9241 | Yes | No | Yes | yes | - | Human, USA | CP009590.1 | Hoffmaster, 2004 |
| FL2013 | Yes (2.5 kb deletion) | No | only partial sequence present | yes | 99.26% identity, 73% coverage | Human, USA | JHQN01000000 | Marston, 2016 |
| BcLA2007 | Yes | No | Yes | No | No pBFH_1 homologues | Human, USA | MUBB00000000 | Pena-Gonzalez, 2017 |
| 03BB87 | No | No | Yes | yes | 100% identity and coverage | Human, USA | CP009941.1 | Johnson, 2015 |
| 03BB102 | Only partial homology | No | No | No | No pBFH_1 homologues | Human, USA | CP001407 | Hoffmaster, 2006 |
| Elc2 | Yes | No | No |  | unknown | Human, USA | ? | Wright, 2011 |
| 03BB108 | Only partial homology. No Anthrax tripartite toxin genes. | No | No | No | No pBFH_1 homologues (pBFI_4: predicted phagemid, 8% coverage. pBFI_5: predicted phagemid, 3% coverage) | Soil, USA | GCA_000832865.1 | Hoffmaster, 2006 |
| BC-AK | Yes | Yes | No | yes | 95.28% identity, 61% coverage | Kangaroo, China Zoo | NZ_CP020937 | Dupke, 2019 |

### *Bc*G9241 sporulates rapidly in comparison to *Bc*ATCC 14579 and shows a temperature dependant phenotype

*B. anthracis* sporulates rapidly under laboratory conditions (Koch, 1876), a phenotype necessary for its obligate infective lifecycle (Ross and Billing, 1957). To determine if the WT strain shares this sporulation phenotype, it was visualised using microscopy after 24 and 48 hours growth on LB agar at both 25 °C and 37 °C. For comparison, *Bc*ATCC 14579 was also imaged under the same conditions (Figure 6A). After 24 hours growth at 25 °C WT cells remained mainly vegetative with little or no endospore formation. However, after 48 hours growth, a limited number of endospores could be observed. In contrast, after 24 hours growth at 37 °C, the WT culture was almost 100% mature spores and we could observe no remaining vegetative cells. Conversely even after 48 hours growth on LB agar at 37 °C, *Bc*ATCC1479 displayed no endospore formation at either temperature. These findings show a striking difference in the sporulation tendency of WT compared to *Bc*ATCC1479, and also that WT sporulated more rapidly at 37 °C compared to 25 °C.

**Figure 6.**
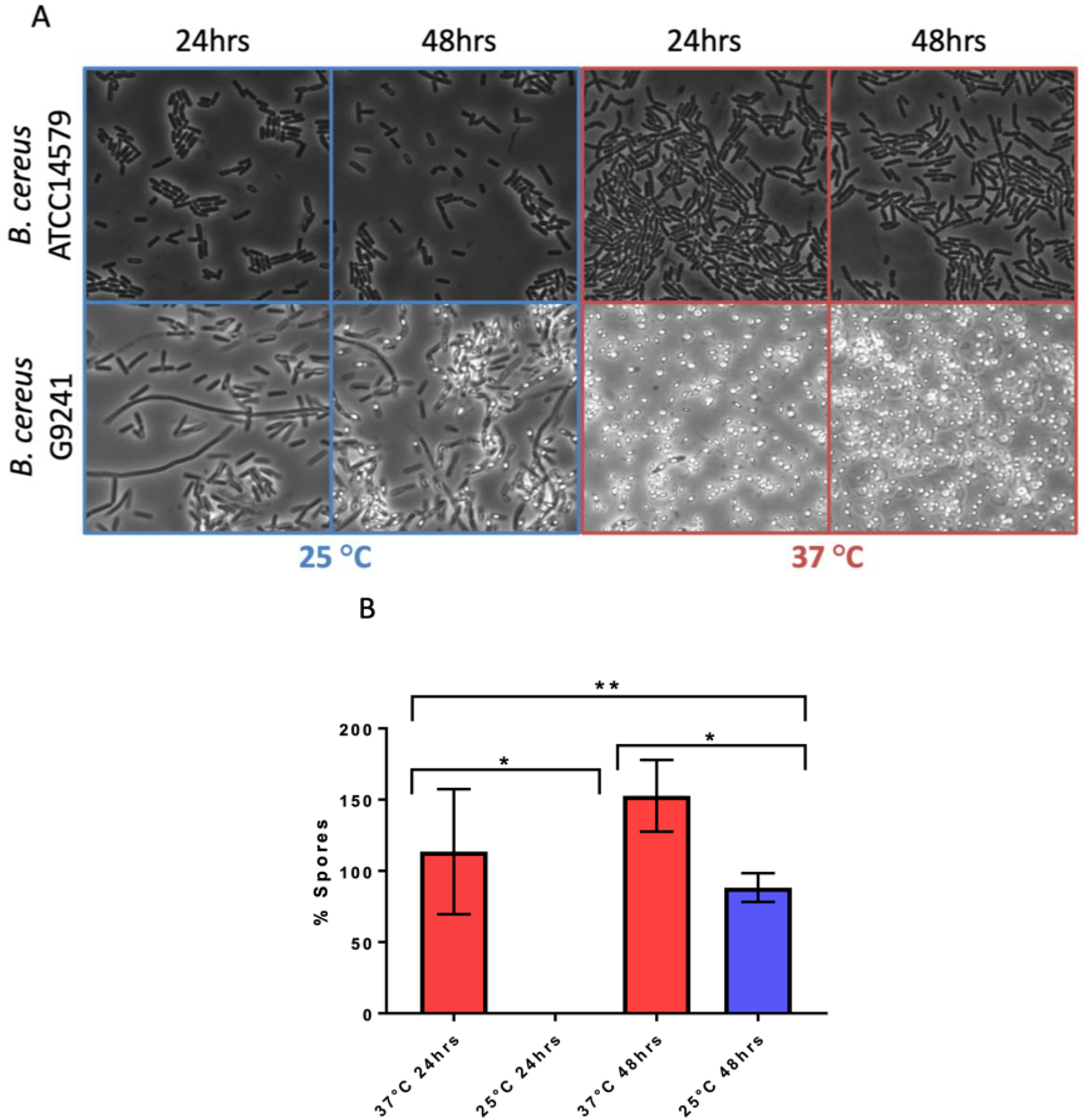
Temperature dependent sporulation of *Bc*G9241 and *Bc*ATCC 14579 in LB. **A**) *Bc*G9241 WT and *Bc*ATCC 14579 were grown for 24 or 48 hrs on LB agar at either 25 °C or 37 °C. A loop of bacterial lawn was taken and used to inoculate 25 μl of PBS. 5 μl of this suspension was mounted and images were taken at 100x magnification. No spore formation is seen under any condition for *Bc*ATCC 14579. While *Bc*G9241 WT had not formed any spores within 24 hrs at 25 °C, by 48 hrs growth endospores have begun to form. In contrast, after only 24 hrs growth at 37 °C, fully mature spores had formed and few if any vegetative cells remained. **B**) *Bc*G9241 WT was cultured in LB broth at 25 °C and 37 °C with aeration. At 24 and 48 hours samples were collected and a sporulation assay was performed to quantify percentage of spores. Data is plotted as the % of the culture that had formed spores. After 24 hours at 37 °C, 100% of the cell population had formed spores. However, at 25 °C, only 0.06% of cells have formed spores. * indicates significance to a p-value < 0.05 using Welch’s t test. **indicates significance to a p-value of 0.0014 using an ordinary one-way ANOVA. Error bars are one standard deviation and all tests are in triplicate (n=3). See text for an explanation of readings of >100 %.

To quantify the temperature dependant sporulation of WT, a sporulation assay was carried out in LB broth at 37 °C and 25 °C. Representative microscopic images were taken to confirm results of the sporulation assay (Figure S3). After 24 hours growth there was a significant difference in percentage of spores between the WT cultures grown at 37 °C and those at 25 °C. After 24 hours at 37 °C, measurements suggested 114% of cells had formed spores, compared to a negligible percentage at 25 °C (Figure 5B). The phenomenon of a >100% spore count (using this assay) has been observed in previous publications (Dale *et al*., 2018), and it is believed to arise from the germination of spores and cell division, after boiling but before plating. At 48 hours there was still a significantly higher percentage of spores formed at 37 °C than at 25 °C. The results of the sporulation assay are consistent with the microscopy analysis of samples (Figure S3). When comparing the proportion of spores formed in liquid and solid media we can see that WT sporulated around 24 hours earlier on the LB agar (Figure 5A, Figure S3).

### pBCX01 is not involved in the rapid sporulation phenotype of *Bc*G9241

A sporulation assay comparing the percentage spore formation of WT, ΔpBCX01 and *Bc*ATCC 14579 at 37 °C and 25 °C was carried out to determine whether the presence of pBCX01 contributes to the rapid sporulation phenotype of *Bc*G9241 when grown under sporulation inducing conditions. Plasmid pBCX01 carries a Rap-Phr system with 100% identity to that encoded on pXO1 from *B. anthracis* (Bongiorni *et al*., 2007). This system is thought to prevent sporulation of *B. anthracis* under infection relevant conditions when optimal toxin production and rapid vegetative growth is advantageous. Therefore, these experiments were carried in the sporulation inducing media, modified G medium (MGM), to allow the effect of pBCX01 to be studied under conditions that don’t inhibit spore formation.

WT, ΔpBCX01 and *Bc*ATCC 14579 were grown in MGM at 37°C for 8 hours and 25°C for 20 hours before the percentage spore formation was assessed. These assays showed that the sporulation phenotype of *Bc*G9241 is not significantly affected by the presence of pBCX01 at either 37 °C or 25 °C (Figure 7). Both WT and ΔpBCX01 form a significantly higher percentage of spores than *Bc*ATCC 14579 at 37 °C or 25 °C at the time points sampled (Figure 7).

**Figure 7.**
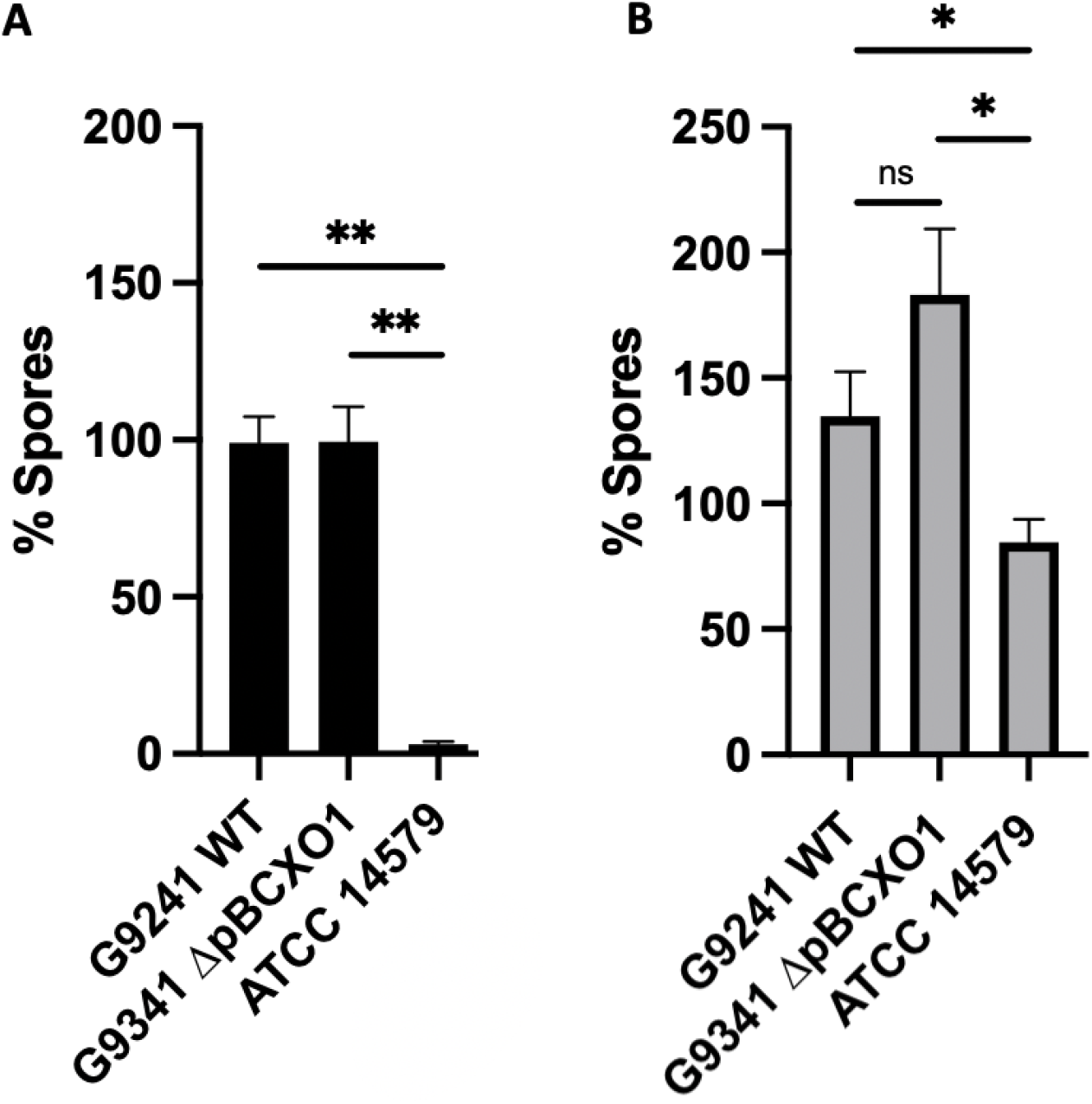
Percentage spore formation in MGM. *B*cG9241 WT, *Bc*G9241 ΔpBCXO1 and *Bc*ATCC 14579 were grown in modified G medium (MGM) for 8 hours at 37°C (**A**) or 20 hours at 25°C (**B**) with shaking at 200 rpm. Percentage of spores in each samples was calculated using a sporulation assay. The black and grey bars show the average percentage of spores from three biological replicates with the standard error of the mean as a line above the bar. The horizontal lines indicate the statistical comparisons made between two samples using an unpaired, Welch’s t-test where ‘not significant’ (ns) = P > 0.05, * = P ≤ 0.05, ** = P ≤ 0.01, *** = P ≤ 0.001.

As *Bc*G9241 sporulates rapidly at 37 °C compared to *Bc*ATCC 14579 we decided to compare the sporulation processes of each strain. We constructed fluorescent reporter strains, designed to express GFP at two critical transcriptional check-points in the sporulation cascade. The aim was to investigate which point in the sporulation cascade is influenced by a temperature difference. Two promoters were selected to create the GFP reporters that would be specifically activated at (i) the initiation of the sporulation cascade and (ii) completion of DNA engulfment into the forespore. These two promoters are activated by the regulators; Spo0A, in its phosphorylated form, (Strauch *et al*., 1992) and Sigma G (Regan *et al*., 2012) respectively. These reporter strains showed that both WT and ΔpBCX01 activated the sporulation cascade at a slightly earlier time point than *Bc*ATCC 14579 and reached the late forespore stage at around 3 hours before *Bc*ATCC 14579 when grown at 37°C in MGM (Figure S4-S5). For brevity the findings of this more detailed analysis have been included in the supplementary data.

## DISCUSSION

An analysis of the influence of both pBCX01 and temperature on the transcriptomic landscape of *Bc*G9241 clearly demonstrates that pBCX01 has a strong influence on the transcriptome of *Bc*G9241 at the mammalian infection relevant temperature of 37 °C. Notably there is significantly less influence of this plasmid at the more environmentally appropriate temperature of 25 °C. Our findings are consistent with experiments in *B. anthracis* where pXO1 has been shown to contribute to both plasmid and chromosomal gene regulation under mammalian infection relevant conditions (McKenzie *et al*., 2014).

Comparing the differential expression at 37 °C and 25 °C between WT and ΔpBCX01, we see that under 37 °C growth conditions removal of pBCX01 had a positive effect on genes one would associate with an increase growth rate in comparison to 25 °C. Unexpectedly, when the plasmid is present (in the WT strain) the same pattern was not seen even though both strains exhibit a similar increase in growth rate at 37 °C compared to 25 °C. This suggests that the rich growth medium used is able to provide sufficient resources to support this increased biomass production irrespective of changes in the transcriptome. The influence of pBCX01 on the transcriptome results in the suppression of transcription of genes involved in various aspects of central metabolism and the biosynthesis of secondary metabolites and amino acids. This effect is likely more important during an actual infection process. Consistent with our findings, previous studies have shown that in *B. anthracis*, AtxA has a negative effect on the expression of amino acid biosynthesis genes (Bourgogne *et al*., 2003; Panda *et al*., 2014). Taken together these results suggest that at 37 °C pBCX01 is having a large effect on gene expression and is acting to suppress the metabolic and biosynthetic processes associated with a higher growth rate. Whereas at 25 °C, the plasmid is having a minimal effect on the transcriptome and there is no evidence of a similar suppression of metabolic and biosynthetic genes we see at 37 °C. Another temperature dependent effect of pBCX01 on the transcriptome appears to be a suppression of the transcription of bacterial chemotaxis genes at 37 °C but not at 25 °C.

Although pBCX01 has a negative effect on the expression of genes encoding amino acid biosynthesis proteins, the WT was still able to synthesize most amino acids not made available in the defined growth media, and indeed even had a growth advantage over ΔpBCX01. It therefore appears that the level of amino acid biosynthesis enzymes in WT, is still sufficient to service the need for biomass production *in vitro*. The suppressive effect of pBCX01 may represent an adaptation to favor amino acid import during a mammalian infection, rather than expend unnecessary energy in *de novo* synthesis. In support of this, we note the expression of amino acid transporters was enhanced by the presence of pBCX01, again suggesting that pBCX01 promotes the uptake of available nutrients in favor of *de novo* biosynthesis.

Another striking effect of the carriage of pBCX01 at 37°C is an increase in the transcription of many diverse transmembrane proteins. Furthermore, both pBCX01 and temperature had an effect on the expression of multiple genes associated with maintenance of the cell membrane of *Bc*G9241. This implies that both the virulence plasmid and temperature both effect how *Bc*G9241 interacts with its environment. We note that pBCX01 encodes the *hasACB* operon for the synthesis of a hyaluronic acid capsule as well as other surface associated proteins such as BslA and an S-layer associated protein, which has been shown to mediate adhesion to host cells (Wang *et al*., 2013; Scarff *et al*., 2018). It is possible that one or more of the uncharacterized transmembrane proteins showing a higher transcript abundance in the WT strain might be involved in the secretion or assembly of the hyaluronic acid capsule or other plasmid encoded surface proteins. Another notable consequence of the carriage of pBCX01 is an increase in transcription of several genes encoding proteins containing helix-turn-helix domains. Indeed, two of these genes were in the top three most differentially expressed genes showing higher levels when pBCX01 is present. This suggests that the presence of pBCX01, and possibly AtxA, can activate the expression of other DNA binding proteins that likely regulate chromosomal genes.

Modulation of metabolism has been shown to be a strategy used by many bacterial pathogens to aid successful infection (Olive and Sassetti, 2016). We propose that the influence of AtxA, or other pBCX01 encoded factors, on the transcription of chromosomal genes may facilitate virulence in *Bc*G9241. This may act by promoting the use of imported host nutrients, while at the same time suppressing unnecessary biosynthesis of required nutrients and metabolites for growth and biomass production. This strategy may give *Bc*G9241 a growth advantage during infection and therefore increasing pathogenic potential. Indeed, higher numbers of tissue resident bacilli has previously been associated with an increased likelihood of fatal outcomes in anthrax infections (Guarner *et al*., 2003). Furthermore, depriving and weakening the host cells of nutrients through increased import has also been proposed to constitute a specific pathogenic strategy in a phenomenon named “nutritional virulence” (Abu Kwaik and Bumann, 2013). Finally, certain metabolic products produced by a pathogen can also have the potential to activate an immune response (Wynosky-Dolfi *et al*., 2014; Gaudet *et al*., 2015). Therefore, the modulation of the metabolism of *Bc*G9241 by pBCX01 may serve to limit the production of potential immune reactive metabolic products.

Sporulation is an essential part of the *B. anthracis* lifestyle and it has been shown to sporulate rapidly under favorable conditions (Davies, 1960). Analysis of *Bc*G9241 sporulation showed that it is also able to sporulate rapidly compared to *Bc*ATCC 14579 in both nutrient rich and nutrient poor sporulation media. This difference is also more pronounced at 37 °C than at 25 °C. Whilst there is a clear difference between the sporulation phenotypes of *Bc*G9241 and *Bc*ATCC 14579, our findings demonstrate that it is not facillitated by AtxA or the presence of pBCX01 as ΔpBCX01 exhibits the same rapid sporulation phenotype. This suggests that there are other genetic elements driving the rapid sporulation phenotype in *Bc*G9241. Interestingly, this work has identified that the pBFH_1 plasmid is highly expressed at 37 °C in comparison to 25 °C and leads to the production of Siphoviridae-like phage particles. There have been multiple examples of bacteriophages enhancing the sporulation efficiency of their host and it is possible that higher expression and production of pBFH_1 proteins at 37°C is influencing the regulation of sporulation (Silver-Mysliwiec and Bramucci, 1990; Peng and Yuan, 2018). As shown in Table 4, pBFH_1-like plasmids are maintained by multiple ‘cross-over’ strains suggesting that it could play an essential role in the biology of a *B. cereus* strain causing an anthrax-like disease.

## AUTHORS AND CONTRIBUTORS

**Grace Taylor-Joyce**^**1**^: Planned and performed experiments and wrote much of the manuscript

**Shathviga Manoharan**^**1**^: Assisted in experiments and wrote parts of the manuscript

**Thomas Brooker**^**1**^: Planned and performed some of the experiments.

**Carmen Sara Hernandez-Rodriguez**^**2**^: Planned and performed some of the experiments.

**Les Baillie**^**3**^: Provided certain bacterial strains and provided advice on handling them.

**Petra Oyston**^**4**^: Provided advice on handling the pathogenic strains and assisted in interpreting the results.

**Alexia Hapeshi**^**1**^: Assisted in some experimental work and in interpreting certain results.

**Nicholas R. Waterfield**^**1**^: Experimental planning, secured funding, assisted in interpreting results and provided guidance and edits for writing the manuscript.

^1^Division of Biomedical Sciences, Warwick Medical School, University of Warwick, Gibbet Hill Road, Coventry, CV4 7AL, United Kingdom

^2^Institut Universitari de Biotecnologia i Biomedicina, Departament de Genètica, Facultad de Ciències Biològiques, University of Valencia, 46100 Burjassot, Valencia, Spain

^3^School of Pharmacy and Pharmaceutical Sciences, Cardiff University, CF10 3AT, Cardiff, United Kingdom

^4^CBR Division, Dstl Porton Down, Salisbury, SP4 0JQ, United Kingdom

## ACKNOWLEDGEMENTS

We would like to thank the Petra Oyston at the DSTL for advice throughout this project.

## MATERIALS AND METHODS

### Bacterial strains and culture conditions

The bacterial strains used in this study were *Bc*G9241 (Hoffmaster *et al*., 2004), *Bc*G9241 ΔpBCX01 and *B. cereus* reference strain ATCC 14579 (American Type Culture Collection, Manassas, Va.). Overnight cultures were grown in 5 ml of Luria Bertani (LB) broth with appropriate antibiotics at either 25 °C or 37 °C. Before *Bacillus* cultures were seeded, pre-cultures were used to synchronise bacterial cell growth by diluting overnight cultures 1 in 100 in 5 ml LB and grown to exponential phase. Pre-cultures were then diluted to an OD_600_ = 0.005 in a specified media and volume.

### RNA extraction for RNAseq

Overnight cultures of *Bc*G9241 and *Bc*G9241 DpBX01 were grown at 25 °C or 37 °C. Cultures were diluted to OD_600_ = 0.05 and incubated at the corresponding temperature until they had grown to OD_600_= 0.5. This pre-culture was diluted to OD_600_ = 0.005 and grown in 50 ml of LB broth. Cells were cultured to mid exponential phase and collected by centrifugation at 10,000g for 1 minute and pellets were resuspended in 5x volume of Qiagen RNA-protect. Resuspended pellets were stored at -20 °C or used immediately. 1 ml of QIAzol (Qiagen) was added to each pellet suspension before being transferred to Lysing Matrix B tubes (MP Biomedicals). Cells were lysed using the FastPrep®-24 Classic instrument with a COOLPREP™ adapter (MP Biomedicals). Bead beating was conducted at 6 ms^-1^ for 40 s for 2 cycles, with a 300 s pause between cycles. Lysates were centrifuged for 1 minute at 10,000 g and supernatant extracted. Cell lysates were processed for total RNA extraction using the RNeasy Micro Kit (Qiagen) and treatment with RNase-free DNase using the Ambion™ DNase I protocol (ThermoFisher Scientific), following the manufacturer’s protocols. Details of RNAseq library preparation, sequencing and analysis methods can be found in supplementary methods.

### Generation of pBCX01 plasmid-cured *Bc*G9241

For the generation of strain *Bc*G9241 ΔpBCX01 (pBCX01^-^/pBC210^+^/pBFH1^+^), a liquid culture of *Bc*G9241 was grown overnight in LB broth at 42 ºC with shaking at 200 rpm. On each of six consecutive days, the culture was diluted 1/1000 into fresh LB broth and grown as above. Dilutions of the final culture were plated onto NBY-Bicarbonate agar (Wilson *et al*., 2011) and growth at 37 ºC in 5 % CO_2_ to promote capsule production. Individual colonies were passaged several times onto fresh NBY-bicarbonate plates and incubated as described above. Since AtxA, positive regulator of the capsule operon (Uchida *et al*., 1997), is encoded in pBCX01 plasmid, pBCX01^-^ mutant colonies show a rough appearance, distinguishable from the smooth colonies producing the capsule. For PCR screening of candidate strains, individual rough colonies were used to conduct colony-PCR using GoTaq Green master mix (Promega) and a primer set unique to different regions on each of the *Bc*G9241 plasmids pBCX01 and pBC210. Control reactions designed to amplify a region of *Bc*G9241 chromosome (*plcR* and ribosomal *16S* genes) and pBFH1 plasmid were included (Table S5). PCR assays were done under the following conditions: an initial step of 10 min at 95 ºC followed by 30 cycles of 95 ºC for 5 min, 45 ºC for 1 min, and 72 ºC for 1.5 min; final extension for 7 min. As a result of this screening *Bc*G9241 ΔpBCX01 was selected for further experiments. This strain was sequenced on an illumina MiSeq platform to confirm the loss of pBCX01 and a lack of any other genomic alterations (not shown).

### Extraction of pBFH1 phage particles and electron microscopy

Exponential pre-cultures of *Bc*G9241 were diluter to OD 0.005 in 50 ml of LB broth and grown to exponential phase at 37 °C. The culture was then spun at 6,000 g for 10 min. The supernatant was removed and filtered through an 100K Amicon Ultra-15 centrifugal filter column (Millipore) by centrifugation at 4,500 g until all supernatant had been filtered. 500 μl of SM buffer (100mM NaCl, 8mM MgSO_4_ and 50mM Tris HCl) was added to the column filter and vortexed vigorously to resuspend any bacteriophage caught by the filter. The sample was removed from the column and filtered through a 0.2 μm filter to remove any bacteria. Samples were stored at 4 °C. To confirm the presence of pBFH_1 phagemid particles, samples were first treated with DNase I (NEB) then heated to 75°C for 10 min to inactivate the DNase. To break open any phage particles, proteinase K was added and incubated at 37°C for 1 hour before boiling at 100°C for 1 hour. The samples were then screened for the presence of phagemid DNA using two sets of primers (pBFH1_5877_Foward, pBFH1_5877_Reverse, pBFH1_5899_Foward and pBFH1_5899_Reverse) each targeting a pBFH_1 phagemid gene. A primer pair targeting the *fusA* gene was also used to control for the presence of non-phage particle DNA. Phage samples were negatively stained using 2% uranyl acetate for 4 minutes. Micrographs were collected on the JEOL2011 electron microscope using a US1000 CCD camera (Gatan Inc) at various levels of magnification.

### Bacterial nutritional studies

Pre-cultures of *Bc*G9241 WT, *Bc*G9241 ΔpBCX01 and *Bc*ATCC 14579 grown in LB broth were first washed in BSM media (Terwilliger *et al*., 2015) (Table S6), where all amino acids of interest had been omitted, three times before diluting to an OD600 = 0.1 in media with the appropriate amino acid omitted. Samples were then aliquoted into a 96 well plate with 100 ml per well with three technical replicates and grown at 37°C with 700rpm shaking. OD600 readings were taken every 15 min for 24 hours. Three biological replicates were carried out for each sample with one biological replicate per plate. All OD600 readings were normalized to a blank media reading and plotted.

### *Bacillus* sporulation assay

All cultures were grown at either 37 °C or 25 °C. Pre-cultures were diluted to an OD = 0.005 into 50 ml LB broth or 20 ml of MGM broth (Table S6) and grown with 200 rpm shaking before samples were taken at specified time points. Samples were diluted and plated on to LB agar to calculate total number of CFU before heat shocking at 65°C for 30 min to kill any *Bc*G9241 vegetative cells and 70 °C for 15 min to kill any *Bc*ATCC 14579 vegetative cells and plating again onto LB agar to calculate number of spores able to form colonies. Percentage of spores was then calculated.

### Microscopy

2 μl of sample was applied to a prepared agarose pad and a cover slip placed over them. Images were captured on a Leica DMi8 premium-class modular research microscope with a Leica EL6000 external light source, using an ORCA-Flash4.0 V2 Digital CMOS Hamamatsu Camera at 100x magnification.

