## Supplementary figures for "The influence of extrachromosomal elements in the anthrax “cross-over” strain *Bacillus cereus* G9241" (from another version of this preprint: version 1, 2022-07-13)

**SUPPLEMENTARY DATA**

Grace Taylor-Joyce[1], Shathviga Manoharan[1], Thomas Brooker[1], Carmen Sara Hernandez-Rodrıguez[2], Les Baillie[3], Petra Oyston[4], Alexia Hapeshi[1], and Nicholas R. Waterfield[1]✉.

[1]Division of Biomedical Sciences, Warwick Medical School, University of Warwick, Gibbet Hill Road, Coventry, CV4 7AL, United Kingdom

[2]Institut Universitari de Biotecnologia i Biomedicina, Departament de Genètica, Facultad de Ciències Biològiques, University of Valencia, 46100 Burjassot, Valencia, Spain

[3]School of Pharmacy and Pharmaceutical Sciences, Cardiff University, CF10 3AT, Cardiff, United Kingdom

[4]CBR Division, Dstl Porton Down, Salisbury, SP4 0JQ, United Kingdom

#### SUPPLEMENTARY RESULTS

**Differential expression datasets.** Raw differential expression data generated by Deseq2 can be found in supplementary csv documents. **Sup_1_DEG_delta37E_WT37E_deseq2.csv** contains differential expression data from the comparison of *Bc*G9241 ΔpBCXO1 and *Bc*G9241 WT grown at 37 ˚C mid-exponential phase, log-fold change values are calculated in respect to the *Bc*G9241 ΔpBCXO1 sample. **Sup_2_DEG_delta25E_WT25E_deseq2.csv** contains differential expression data from the comparison of *Bc*G9241 ΔpBCXO1 and *Bc*G9241 WT grown at 25 ˚C mid-exponential phase, log-fold change values are calculated in respect to the *Bc*G9241 ΔpBCXO1 sample. **Sup_3_DEG_WT25E_WT37E_deseq2.csv** contains differential expression data from the comparison of growth at 25 ˚C and 37 ˚C of *Bc*G9241 WT mid-exponential phase cultures, log-fold change values are calculated in respect to 25 ˚C grown samples. **Sup_4_DEG_delta25E_delta37E_deseq2.csv** contains differential expression data from the comparison of growth at 25 ˚C and 37 ˚C of *Bc*G9241 ΔpBCXO1 mid-exponential phase cultures, Log-fold change values are calculated in respect to 25 ˚C grown samples.

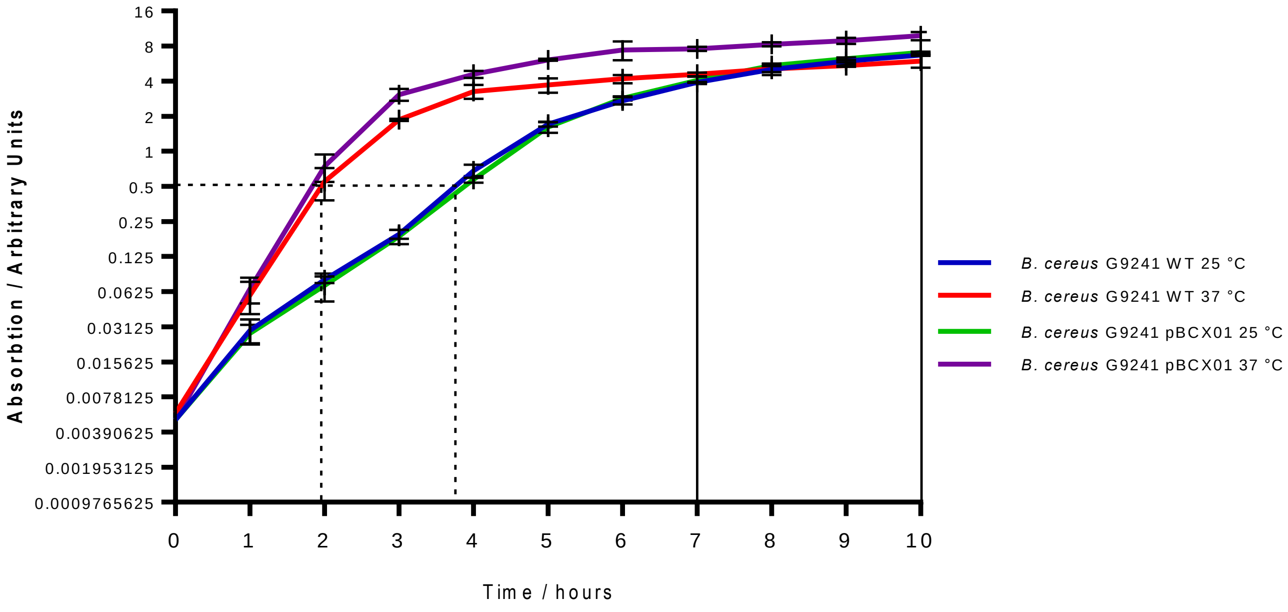


**Figure S1. growth rate of *Bc*G9241 WT and *Bc*G9241 pBCX01 at 25 °C and 37 °C**. Strains were grown in LB broth for 10 hours, shaking at 200 rpm. A pre-culture step was used to remove a lag phase. *Bc*G9241 WT and ΔpBCX01 grow almost identically at 25 °C (blue and green lines respectively). However, at 37 °C the WT (red line) grows slower than the ΔpBCX01 strain (purple line). Dashed lines indicate time points for mid-exponential phase ($OD_{600}$ = 0.5). Solid black lines indicate stationary phase time points, estimated to be 4 hours after transition phase (not annotated). All points plotted are averages of at least 3 measurements (n=3).

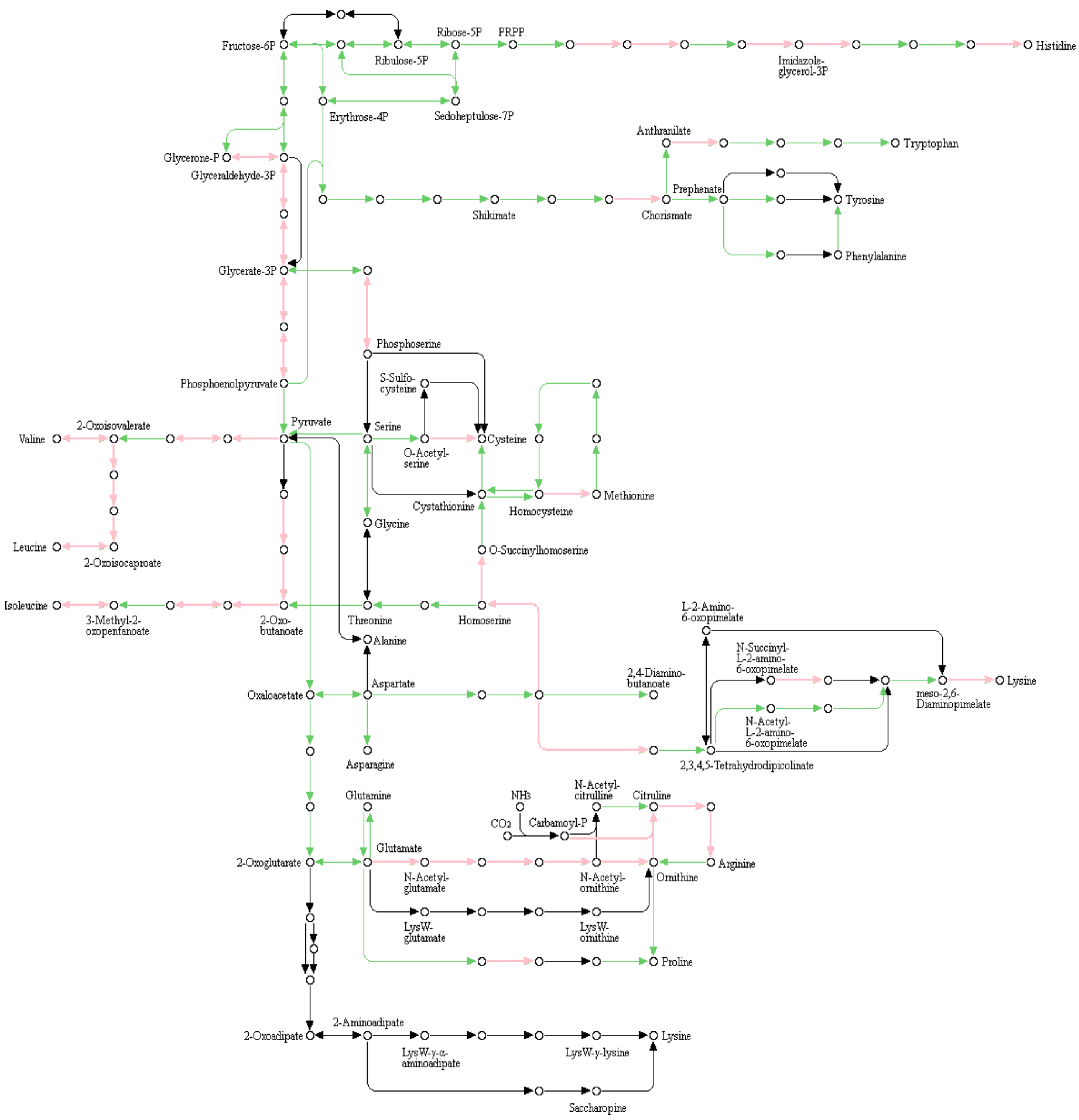


**Figure S2: Map of AA biosynthetic pathways generated by KEGG.** Arrows represent proteins and dots represent the products synthesized by the proteins or the substrate of the protein. Green and pink arrows indicate proteins encoded by genes present in *B. anthracis* Sterne. Pink genes also represent genes that were more highly expressed in *Bc*G9241 ΔpBCX01 compared to WT when grown at 37˚C in LB at either exponential or stationary phase. KEGG reference: Kanehisa, M. and Sato, Y.; KEGG Mapper for inferring cellular functions from protein sequences. Protein Sci. 29, 28-35 (2020). [pubmed] [pdf]

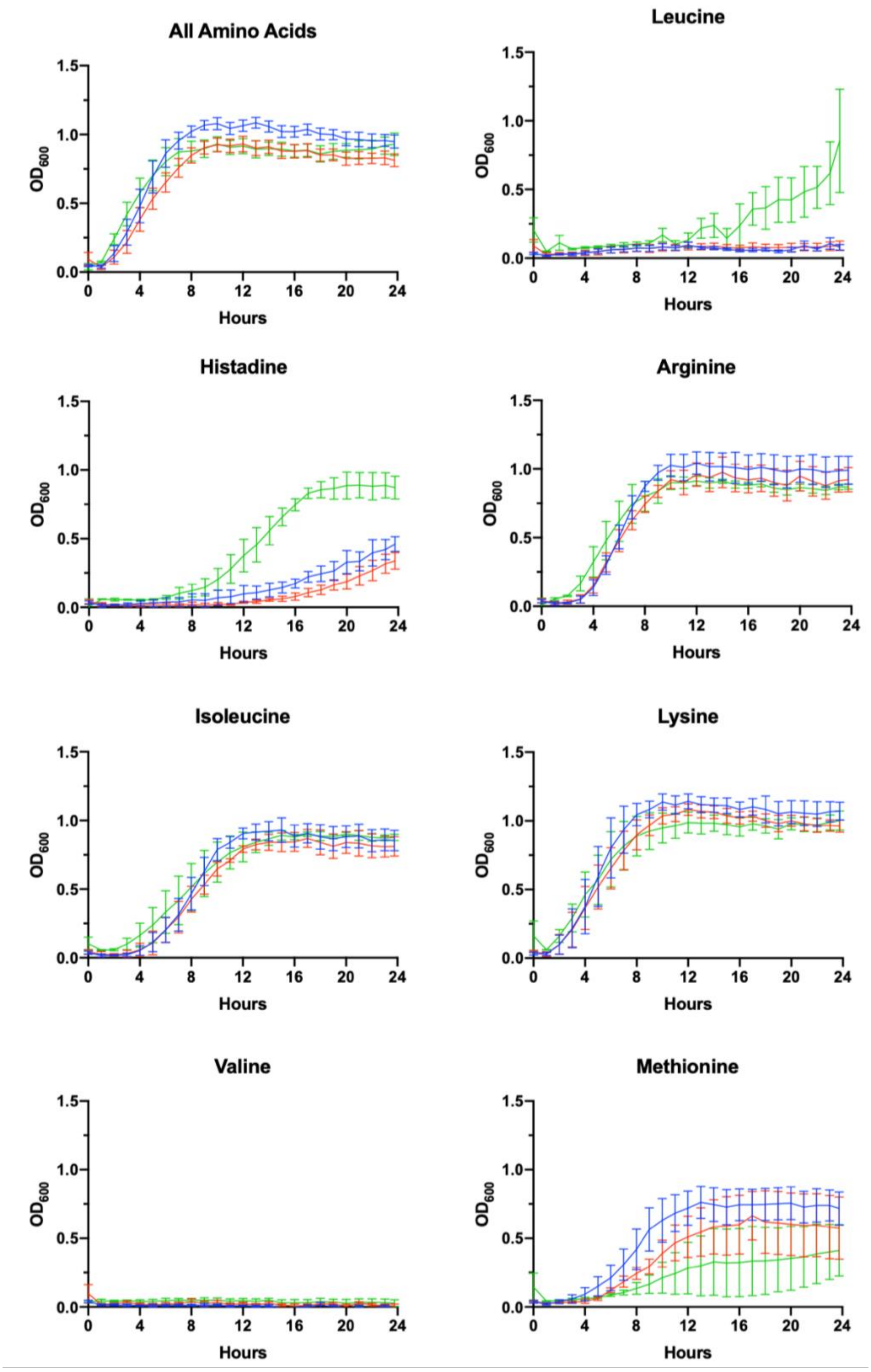


**Figure S3: Growth in AA limiting media.** Growth curves showing optical density measured at 600mn wavelength ($OD_{600}$) plotted against time of *Bc*G9241 WT (blue line), *Bc*G9241 ΔpBCX01 (red line) and *Bc*ATCC 14579 (green line). The bars show standard error. Cultures were grown at 37˚C with shaking in blood serum mimic (BSM) media with a certain amino acid omitted.

The amino acid written above each graph indicates which amino acid was omitted from the growth media.

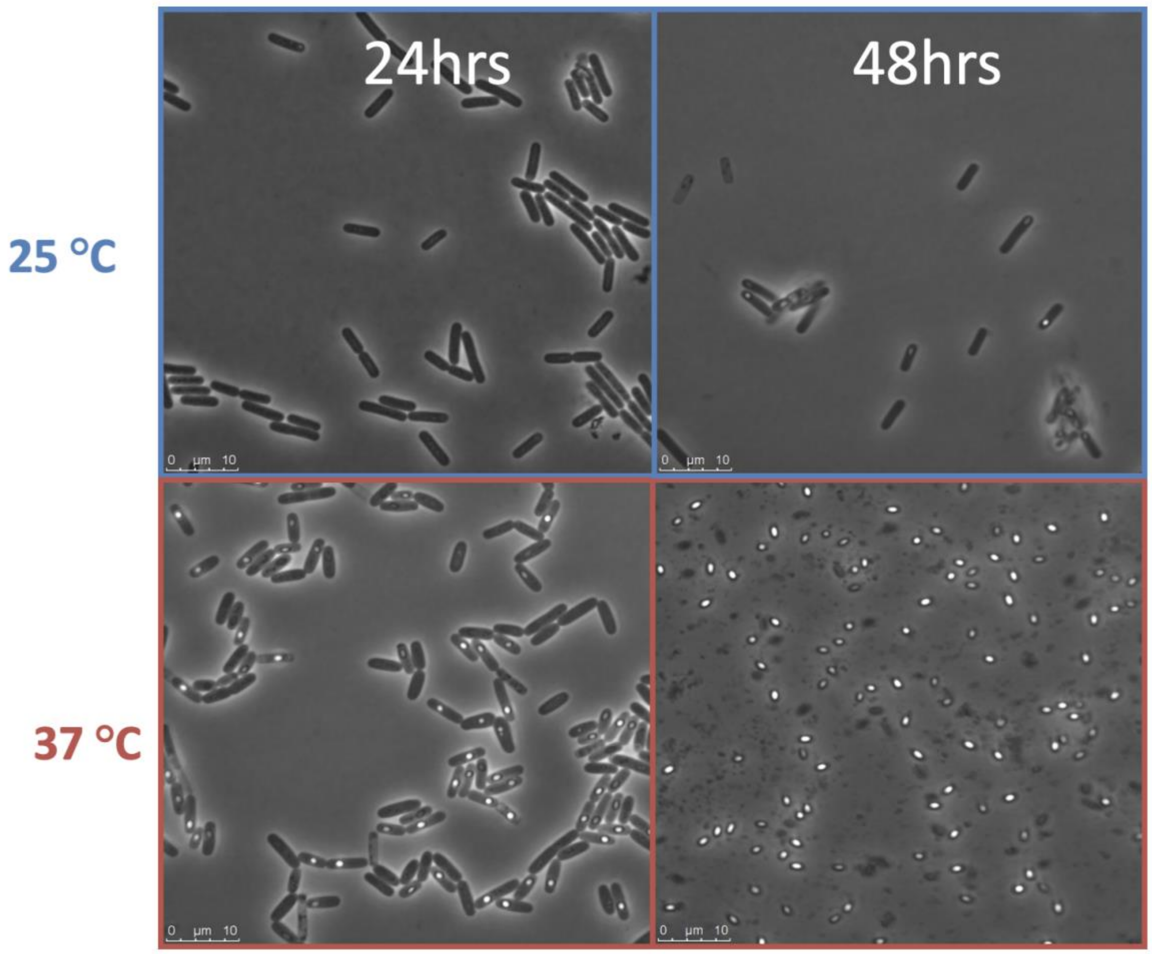


**Figure S4. Microscopic examination of *Bc*G9241 WT grown at 25 °C and 37 °C in LB broth.** *Bc*G9241 WT was grown at 25 °C and 37 °C in LB broth with aeration for up to 48 hrs. Aliquots of 2 µl of bacterial cultures were pipetted onto agarose pads and imaged at 100x magnification after 24 and 48 hrs. After 48 hrs at 25 °C, *Bc*G9241 cell populations were observed to be heterogeneous, with some cells containing endospores, and others apparently remaining vegetative. However when grown at 37 °C, *Bc*G9241 sporulated more rapidly, with the majority of cells containing endospores by 24 hrs, and a homogeneous population of mature spores at 48 hrs. Spores can be seen as phase bright and vegetative cells as phase dark. Dark specs in the bottom right panel are likely pieces of lysed mother cell debris.

**Analysis of the kinetics of temperature dependent sporulation.**

The promoters from the genes (i) *spo0A* itself and (ii) *sspA* were cloned into a high copy number plasmid upstream of a *gfp* gene before transformation into *Bc*G9241 and *Bc*ATCC14579. Fragments used for the reporter constructs included the transcription promoters and native Shine Dalgarno sequences of the target genes to encompass both transcription and translation activities. The fluorescence of the Spo0A and Sigma G sporulation reporters in WT, ΔpBCX01 and *Bc*ATCC14579 were measured across the growth curve in a microplate reader at 37 ˚C in MGM broth. Fluorescent microscopy images of both reporters cultured at 37˚C in MGM broth (Figure S5-S6).

Fluorescent readings taken over time at 37 ˚C of the Spo0A reporter strains show an increase in Spo0A driven fluorescence of *Bc*G9241 WT and *Bc*G9241 ΔpBCX01 was observed at around 4 hours, while for *Bc*ATCC14579 it was slightly later at around 4.5 hours (Figure S5A). At 37 ˚C the micrographs of Spo0A reporters (Figure S5B) of *Bc*G9241 WT and *Bc*G9241 ΔpBCX01 at 8 hours and 10 hours show that most cells have visible phase bright forespores and there is evidence of heterogenous activity of Spo0A, where some cells have a higher intensity of fluorescence than others. *Bc*G9241 WT and *Bc*G9241 ΔpBCX01 Spo0A reporters have the same phenotype at all time points observed. At 8 hours over half of *Bc*ATCC14579 cells contain a forespore and by 10 hours nearly all cells have a visible forespore, similar to *Bc*G9241. At 8 hours and 10 hours there are a small number of *B*cATCC14579 cells that are florescent, fewer than *Bc*G9241 WT or *Bc*G9241 ΔpBCX01 reporters and with lower florescent intensity. At the later time point of 12 hours the proportion and intensity of florescent *Bc*ATCC14579 cells does not increase (data not shown). This suggests that *Bc*ATCC14579 has less Spo0A activity under these conditions.

Fluorescent readings of the Sigma G reporter strains showed an increase in the Sigma G dependant fluorescence of *Bc*G9241 WT and *Bc*G9241 ΔpBCX01 started around 5 hours, while that of *Bc*ATCC14579 took longer, arising around 9 hours during culture at 37 ˚C (Figure S6A). We note there is a bigger difference between the timing of peak fluorescence of the *Bc*G9241 stains and *Bc*ATCC14579 in the Sigma G reporter strain than the Spo0A reporters (Figure S5-S6).

Micrographs of Sigma G reporter strains showed that at 37 ˚C *Bc*G9241 WT and *Bc*G9241 ΔpBCX01 Sigma G reporters have the same phenotype at all time points observed (Figure S6B). At 8 hours, 37 ˚C, most cells have visible phase bright forespores and florescence localized to the forespore in *Bc*G9241 WT and *Bc*G9241 ΔpBCX01 whereas there is only a small number visible phase bright forespores in *Bc*ATCC14579 and some cells with florescence localized to both early forespores and those that are phase bright (Figure S6B). At 10 hours all living cells have florescent phase bright forespores in *Bc*G9241 WT and *Bc*G9241 ΔpBCX01. At 10 hours most, but not all, *Bc*ATCC14579 cells have florescent phase bright forespores.

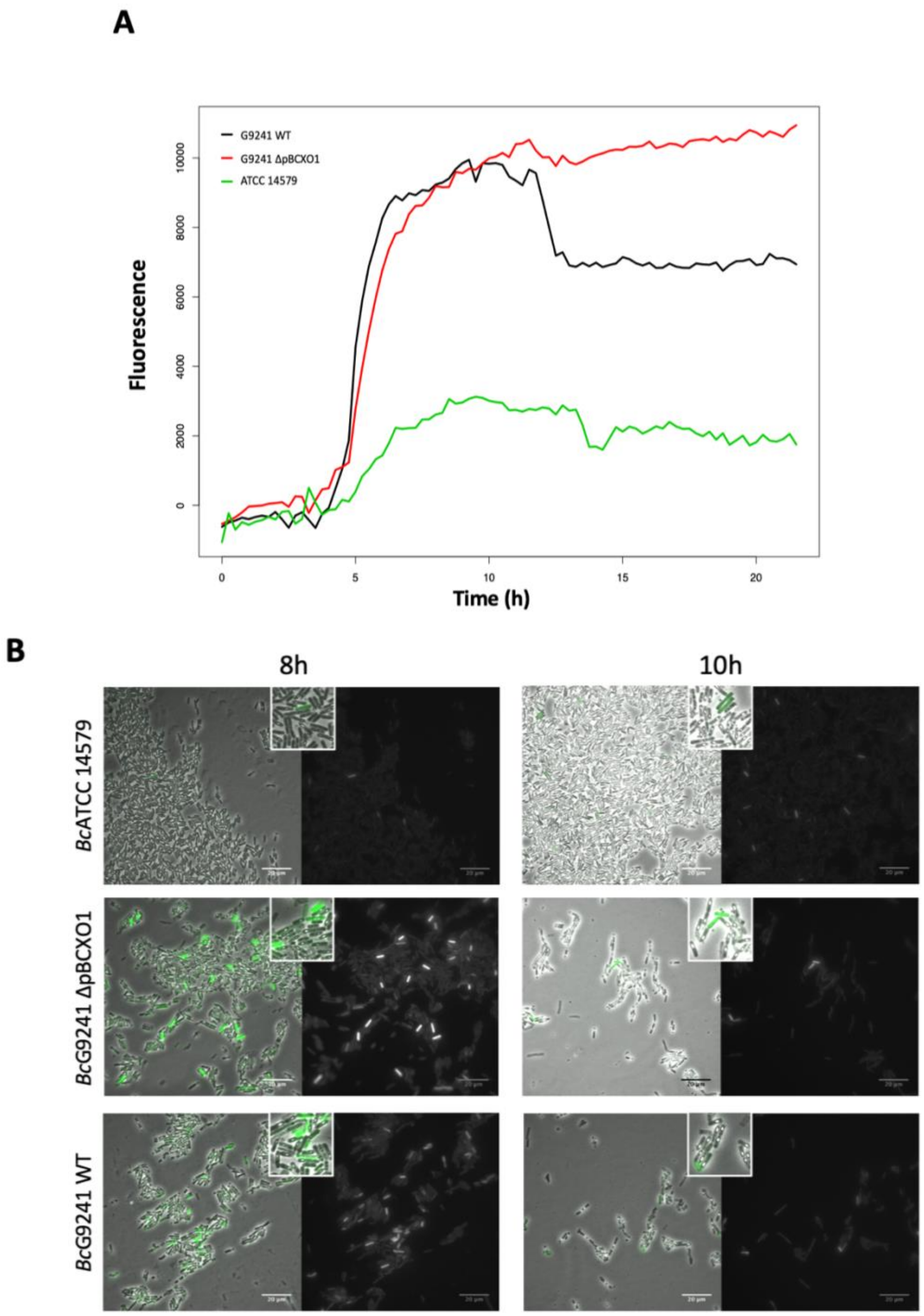


**Figure S5: Activity of Spo0A over time in MGM at 37 ˚C. (A)** Shows the florescence of Spo0A reporter strains of *Bc*G9241 WT, *Bc*G9241 ΔpBCXO1 and *Bc*ATCC14579 grown over 24 hours in 100 µl volume of MGM. Each line represents the mean of three biological replicates with three technical replicates each. (**B**) The *Bc*G9241 WT, *Bc*G9241 ΔpBCXO1 and *Bc*ATCC14579 Spo0A fluorescent reporter strains were grown in 20 mL MGM media and visualizing by fluorescent microscopy at X100 magnification. Grown at 37˚C for 8 and 10 hours. The left image shows the fluorescent and phase contrast channels overlaid and the right image shows only the fluorescence channel.

A

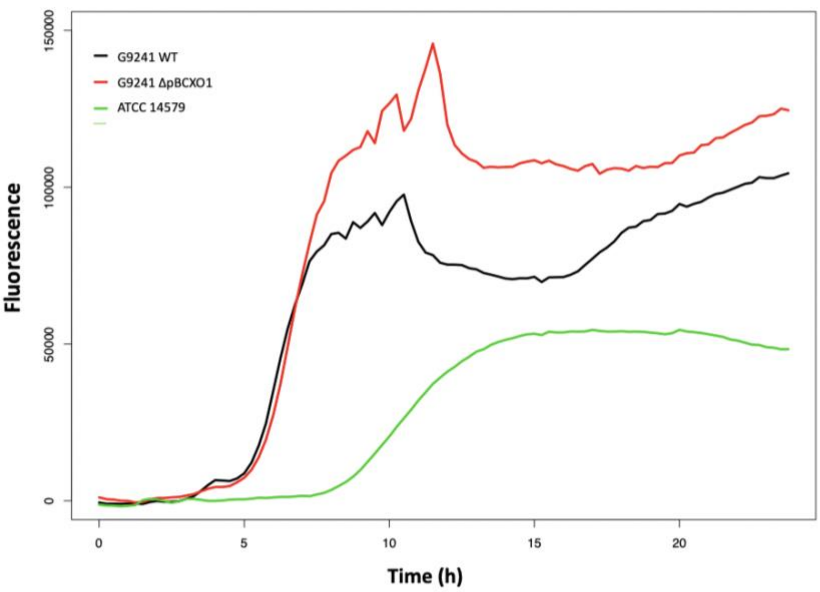


B

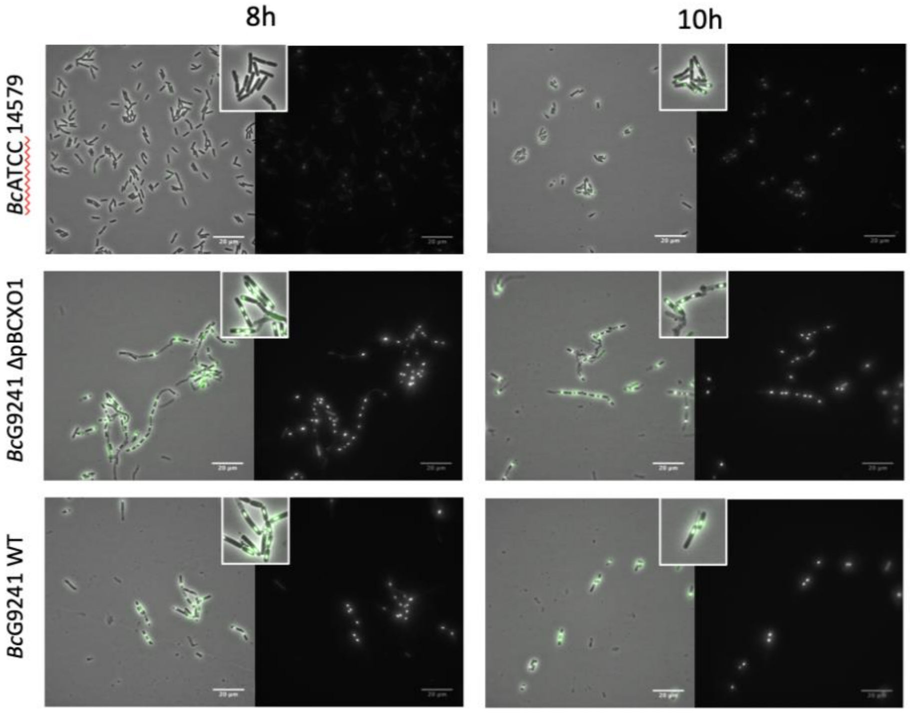


**Figure S6: Activity of Sigma G over time in MGM at 37 ˚C. (A)** Shows the florescence of Sigma G reporter strains of *Bc*G9241 WT, *Bc*G9241 ΔpBCXO1 and *Bc*ATCC14579 grown over 24 hours in 100 µl volume of MGM. Each line represents the mean of three biological replicates with three technical replicates each. (**B**) The *Bc*G9241 WT, *Bc*G9241 ΔpBCXO1 and *Bc*ATCC14579 Sigma G fluorescent reporter strains were grown in 20 mL MGM media and visualizing by fluorescent microscopy at X100 magnification. Grown at 37˚C for 8 and 10 hours. The left image shows the fluorescent and phase contrast channels overlaid and the right image shows only the fluorescence channel.

**Table S1.** The top 15 differentially expressed genes with higher transcription at 37 ˚C compared to 25 ˚C in *Bc*G9241 WT during exponential phase growth with a p value <=0.05.

| **Log2-fold Change** | **37 ˚C v 25 ˚C at exponential phase, in *Bc*G9241 WT** | **Gene** | **Gene Loci (AQ16_)** |
|---|---|---|---|
| 5.26 | TQO small subunit DoxD family protein | | 2331 |
| 4.93 | L-lactate dehydrogenase | | 2981 |
| 4.65 | lactate utilization protein A | *lutA* | 1186 |
| 4.35 | aldehyde dehydrogenase family protein | | 3652 |
| 4.18 | hypothetical protein | | 5820 |
| 4.14 | ATPase subunit of terminase family protein | | 5898 |
| 3.94 | L-lactate dehydrogenase | | 3111 |
| 3.86 | phage terminase%2C large subunit%2C PBSX family | | 5899 |
| 3.80 | phage family protein | | 5822 |
| 3.78 | lactate utilization protein C | *lutC* | 1184 |
| 3.76 | phage tail family protein | | 5835 |
| 3.76 | iron-sulfur cluster-binding protein | | 1185 |
| 3.75 | hypothetical protein | | 5821 |
| 3.74 | putative membrane protein | | 5895 |
| 3.71 | formate acetyltransferase | *pflB* | 2025 |

**Table S2.** The top 15 differentially expressed genes with higher transcription at 37 ˚C compared to 25 ˚C in *Bc*G9241 ΔpBCX01 during exponential phase growth with a p value <=0.05.

| **Log2-fold Change** | **37 ˚C v 25 ˚C at exponential phase, in *Bc*G9241 ΔpBCX01** | **Gene** | **Gene Loci (AQ16_)** |
|---|---|---|---|
| **6.56** | hypothetical protein | | 5876 |
| **6.37** | aldehyde dehydrogenase family protein | | 3652 |
| **6.13** | hypothetical protein | | 5872 |
| **6.04** | L-lactate dehydrogenase | | 2981 |
| **5.82** | TQO small subunit DoxD family protein | | 2331 |
| **5.63** | minor capsid family protein | | 5827 |
| **5.62** | hypothetical protein | | 5820 |
| **5.48** | minor capsid family protein | | 5828 |
| **5.43** | L-lactate dehydrogenase | | 3111 |
| **5.42** | hypothetical protein | | 5869 |
| **5.38** | formate acetyltransferase | *pflB* | 2025 |
| **5.20** | putative phage protein | | 5873 |
| **5.19** | hypothetical protein | | 5879 |
| **5.17** | hypothetical protein | | 5878 |
| **5.16** | dUTPase family protein | | 5883 |

**Table S3.** Primers used in this study

| Primer Name | Primer sequence 5' to 3' |
|---|---|
| **gfp gene primers** | |
| GFP_KpnI_Foward | GATATAggtaccGCTAGCAAAGGAGAA |
| GFP_EcoRI_Reverse | CCGAGCTCgaattcATTATTTGTAGA |
| **Sporulation reporter primers** | |
| spo0Ap_Foward | AATctgcagTGGAATGTGTTGCGAAAGG |
| spo0Ap_Reverse | ATTggtaccCACAGCTTTTCCTCCCTTAC |
| sspAp_Foward | ATCctgcagATATCGCTCTTAGTTTAGCC |
| sspAp_Reverse | ATAggtaccAGAATTTTGGTTTGCCATAGG |
| **pBFH1 screening primers** | |
| pBFH1_5877_Foward | ATGATAACTCTCGCTGAACGGAACGG |
| pBFH1_5877_Reverse | CATCTGAATCAATCTTTCTATTAATCTGTCTTGCTTGTCC |
| pBFH1_5899_Foward | GTGTGTTAAACGCTTTGAAATGCGTTATACGG |
| pBFH1_5899_Reverse | ATGGTTGTATTTATCGATTGGCGTACCG |
| **Confirm generation of BcG9241 ∆pBCX01** | |
| pBCX01_3-Forward | CATGATGACACCACCAATAA |
| pBCX01_3-Reverse | TCACACATACACCAGCAACT |
| pBCX01_51-Forward | AGGTAGGGATTTTTCTGAGG |
| pBCX01_51-Reverse | GTATGGTGGTTCTGCATCTT |
| pBC210_157-Forward | AATTTACCAACAGTCGTTTGAG |
| pBC210_157-Reverse | ATTTGTCCATGTTTTTCATAGC |
| pBC210_69-Forward | AAACAAAGGATAAGGGGAAG |
| pBC210_69-Reverse | TCCGTAATAAATGGTGGTTC |
| pBFH_1_20-Forward | CTACCGAACCAAAACAAGAG |
| pBFH_1_20-Reverse | CCATTACCTACATCGAAAGC |
| *Bc* G9241_plcR-Forward | ATGCACGCAGAAAAAATTAGG |
| *Bc* G9241_plcR-Re*verse* | CTCGCCCTTTTGATAATACA |
| 16S_B27-Foward[a] | AGAGTTTGATCCTGGCTCAG |
| 16S_U1492-Reverse[a] | GGTTACCTTGTTACGACTT |

**Table S4.** Media used in this study

| Name | Formula | Source |
|---|---|---|
| Modified G Medium (MGM) | yeast extract 0.2% w/v | Kim and Goepfert 1974 |
| | CaCl2.2H2O 0.0025% w/v | |
| | K2HPO4 0.05% w/v | |
| | MgSO4.7H2O 0.02% w/v | |
| | MnSO4.4H2O 0.005% w/v | |
| | ZnSO4.5H2O 0.0005% w/v | |
| | CuSO4.5H2O 0.0005% w/v | |
| | FeSO4.7H2O 0.00005% w/v | |
| | (NH4)2SO4 0.2% w/v | |
| Modified Blood Serum Mimic (BSM) media | KPO4 buffer 5mM | Modified from Terwilliger, Swick et al. 2015 |
| | NaCl 140mM | |
| | CaCl2 2.4mM | |
| | MgCl2 0.8mM | |
| | ZnCl2 0.0146mM | |
| | glucose 6mM | |
| | alanine 1.48mM | |
| | arginine 0.156mM | |
| | asparagine 0.352mM | |
| | aspartic acid 0.012mM | |
| | cystine 0.192mM | |
| | glutamic acid 0.232mM | |
| | glutamine 2.08mM | |
| | glycine 1mM | |
| | histidine 0.292mM | |
| | Isoleucine 0.284mM | |
| | leucine 0.472mM | |
| | lysine 0.74mM | |
| | methionine 0.092mM | |
| | phenylalanine 0.218mM | |
| | proline 0.94mM | |
| | serine 0.392mM | |
| | threonine 0.664mM | |
| | tryptophan 0.212mM | |
| | tyrosine 0.24mM | |
| | valine 0.92mM | |
| | biotin 0.00004mM | |
| | hiamine 0.056mM | |

### SUPPLEMENTARY METHODS

#### rRNA removal

After RNA was extracted from *Bc*G9241 cultures for RNAseq, rRNA was depleted from samples using the Ribo-Zero® rRNA removal kit (illumina®) following manufacturer's instructions detailed in the Illumina Ribo-Zero® rRNA removal kit Reference guide (Document # 15066012 v02). The individual wash protocol was followed included the use of the RiboGuard RNase Inhibitor and a total of 2 µg of RNA was used per sample. Following the probe hybridization and rRNA removal steps the sample was cleaned up using ethanol precipitation following the reference guide instructions. rRNA removal was then validated using the Agilent RNA 6000 pico kit (Agilent) as per manufacturer's instructions.

#### RNAseq library prep

TruSeq library prep was conducted using the Low Sample (LS) protocol and manufacturer's instructions were followed which are detailed in the LS protocol of the TruSeq® Stranded mRNA Sample Preparation Guide (Part # 15031047 Rev. E). The first five steps of the 'purify and fragment mRNA' section were skipped and the protocol was started at the 'incubate RFP' step once 13 µl of the Fragment, Prime, Finish Mix had been added to each 5 µl samples if rRNA depleted RNA. From this point the protocol was followed and only modified by adapting it to use 1.5 ml tubes (or PCR tubes where necessary) instead of 96-well plates. In the 'Synthesize Second Strand cDNA' section the End Repair Control was used. In the 'Ligate Adapters' section, adapter tubes were used instead of a plate. Once all steps were completed up to and including the 'Enrich DNA Fragments' section of the LS protocol the library quality and fragment length was analysed using the Agilent high sensitivity DNA Kit on an Agilent 2100 Bioanalyzer instrument as per manufacturer's instructions.

#### Pooling cDNA libraries

To calculate the molarity of each cDNA library sample, based on the average determined fragment length and the concentration of cDNA, the following equation was used:

$$\frac{\text{concentration of library (ng}\mu\text{l}^{-1}\text{)}}{660\ (\text{g mol}^{-1}\text{bp}^{-1})\times\text{average insert size (bp)}} \times 10^6 = \text{Concentration (nM)}$$

cDNA libraries were then diluted to a concentration of 4 nM. Equal volumes of each samples were then combined into pools to give a 4 nM cDNA library pool. 5 µl of the cDNA library pool was then combined with 5 µl of 0.2 M NaOH. This was then vortexed and centrifuged for 1 min at 280 g before incubating at room temperature for 5 min. The 2 nM pool was then diluted to a final concentration of 9.5 pM in pre chilled HTI buffer (supplied with IlluminaTM MiSeq cartridge) at a volume of 630 µl. This sample was used to load the MiSeq cartridge and ran on the Illumina MiSeq following a paired-end protocol as per manufacturer's instructions.

**Generating differential gene expression data**

Bowtie2-build (Johns Hopkins university) was used to build an indexed reference genomes where pBCX01 loci had been removed for samples of *Bc*G9241 ∆pBCX01. The seqtk tool (GitHub) was then used to flip the first read of paired-end sequences before Bowtie2 was used to map reads to the indexed reference genome. The sequence alignment/map (SAM) files generated by Bowtie2 were converted to BAM files using Samtools (GitHub). The BAM files were then indexed and sorted by genome position using Samtools. CoverageBed (Bedtools) was then used to generate count data from the sorted BAM files. The count data (number of mapped sequence reads for each locus) was used as inputted into the R studio package, DESeq2, which was used to generate differential expression data between all sample comparisons.

**Enrichment analysis using STRING**

Firstly, Roary (Github) was used to assign *B. anthracis* Sterne gene locus tags to *Bc*G9241 genes where there was a minimum percentage similarity of 90%. List of differentially expressed genes (DEG) were generated where the log2-fold change observed was 1 or above and the p-adjusted value was 0.05 or below. A list of DEG, using the corresponding *B. anthracis* Sterne gene locus tag, were entered into STRING using the multiple proteins search tool (Szklarczyk et al., 2019). The enrichments in KEGG pathways generated, where there was a false discovery rate of 0.01 or below, were used in this analysis.

**Construction of sporulation reporter vectors.** pHT315 vectors (Arantes and Lereclus, 1991) carrying the *gfp* gene at the multiple cloning site were used to construct the pHT315_spo0Ap_gfp and pHT315_sspAp_gfp sporulation reporter vectors. pHT315_spo0Ap_gfp carries the region of the *spo0A* promoter activated by phosphorylated Spo0A directly upstream of *gfp*. pHT315_sspAp_gfp carries the sspA promoter, which is activated by Sigma G, directly upstream of *gfp*. Vectors were linearized using KpnI and PstI restriction enzymes (NEB). Insert fragments were amplified with Q5 DNA polymerase (NEB) by PCR with the appropriate primer pairs (Table S3) from *Bc*G9241 whole genome extract. The resulting fragments were digested with KpnI and PstI restriction enzymes and ligated into the linearized vector. Plasmid constructs were transformed in to chemically competent *E. coli* DH5-α cells through heat shock. Once confirmed by DNA sequencing, all vectors were transformed into the non-methylating *E. coli* ET12567 strain by electroporation. Vectors amplified by *E. coli* ET12567 were purified and transformed into *B. cereus* strains using electroporation.

**Transformation of *Bacillus cereus* species through electroporation.**
Overnight cultures were diluted 1:100 in 50 ml of LB broth. Cells were cultured at 37 °C and harvested at $OD_{600}$ of 0.5 by centrifugation at 5000 g for 10 minutes at 4˚C. Cell pellets were resuspended in 10 ml of 10% glycerol in $ddH_2O$. The wash was repeated two more times. Washed cell pellets were resuspended in 100 µl of $ddH_2O$ to create ~220 µl of competent cells. 2 µg of plasmid DNA was added to the cells and 50 µl of this cell-plasmid mix was aliquoted into a pre-chilled 2 mm cuvette. Cells were electroporated at 2.5 kV, 25 µF and 200 Ω. 1 ml of SOC media (NEB) was immediately added to aid cell recovery. Cells were transferred to a 15 ml falcon tube and incubated at 37 °C for 3 hours. Transformants were selected on LB agar containing 25 µg/ml of erythromycin.

**Fluorescence plate reader assays of sporulation reporters.** Pre-cultures were diluted to an OD600 = 0.005 in MGM media and 100ul of each sample were aliquoted into a 96 well plate with three technical replicates and three biological replicates. The plates were at either 37˚C or 25˚C with shaking at 700 rpm and fluorescence was measured every 15 min. Each plate contained BcG9241 WT, ∆pBCXO1 and BcATCC14579 strains carrying a fluorescent reporter plasmid with gfp under the control of a promoter of interest as well as each strain carrying a control plasmid with no promoter upstream of gfp. The fluorescence of blank media was subtracted from the readings of all samples and the fluorescence readings of a strain carrying the pHT315-gfp plasmid with no promotor region was used as a fluorescence control strain. Curves showing fluorescence of the whole culture were calculated by subtracting the fluorescent readings of the control strain. Curves showing the fluorescence per cell were calculated by dividing by the fluorescent readings of the control strain.
